# Maximally selective single cell target for circuit control in epilepsy

**DOI:** 10.1101/2020.10.20.340364

**Authors:** Darian Hadjiabadi, Matthew Lovett-Barron, Ivan Raikov, Fraser Sparks, Zhenrui Liao, Scott C. Baraban, Jure Leskovec, Attila Losonczy, Karl Deisseroth, Ivan Soltesz

**Affiliations:** Department of Bioengineering, Stanford University, Stanford, CA, USA; Department of Neurosurgery, Stanford University, Stanford, CA, USA; Neurobiology Section, Division of Biological Sciences, University of California San Diego, La Jolla, CA, USA; Department of Neuroscience, Columbia University, New York, NY, USA; Mortimer B. Zuckerman Mind Brain Behavior Institute, Columbia University, New York, NY, USA; The Kavli Institute for Brain Science, Columbia University, New York, NY, USA; Department of Neurological Sciences, University of California San Francisco, San Francisco, CA, USA; Department of Computer Science, Stanford University, Stanford, CA, USA; Howard Hughes Medical Institute, Stanford University, Stanford, CA, USA

**Keywords:** epilepsy, calcium imaging, effective connectivity, hub neurons, motifs

## Abstract

Neurological and psychiatric disorders are associated with pathological neural dynamics. The fundamental connectivity patterns of cell-cell communication networks that enable pathological dynamics to emerge remain unknown. We studied epileptic circuits using a newly developed integrated computational pipeline applied to cellular resolution functional imaging data. Control and preseizure neural dynamics in larval zebrafish and in chronically epileptic mice were captured using large-scale cellular-resolution calcium imaging. Biologically constrained effective connectivity modeling extracted the underlying cell-cell communication network. Novel analysis of the higher-order network structure revealed the existence of ‘superhub’ cells that are unusually richly connected to the rest of the network through feedforward motifs. Instability in epileptic networks was causally linked to superhubs whose involvement in feedforward motifs critically enhanced downstream excitation. Disconnecting individual superhubs was significantly more effective in stabilizing epileptic networks compared to disconnecting hub cells defined traditionally by connection count. Collectively, these results predict a new, maximally selective and minimally invasive cellular target for seizure control.

**Highlights:**

- Higher-order connectivity patterns of large-scale neuronal communication networks were studied in zebrafish and mice
- Control and epileptic networks were modeled from *in vivo* cellular resolution calcium imaging data
- Rare ‘superhub’ cells unusually richly connected to the rest of the network through higher-order feedforward motifs were identified
- Disconnecting single superhub neurons more effectively stabilized epileptic networks than targeting conventional hub cells defined by high connection count.
- These data predict a maximally selective novel single cell target for minimally invasive seizure control

## Introduction

Methods in network science have been instrumental in deciphering neural communication associated with neurological disorders^1,2^. One such disorder, epilepsy, is characterized by spontaneous and recurrent seizures that arise from abnormal neural activity and synchronization across brain regions^3,4^. Epilepsy impacts over 60 million individuals worldwide and many of these children and adults have medically uncontrolled seizures and suffer from debilitating cognitive and emotional comorbidities. One primary reason for this is that current anti-epileptic treatments lack spatial, temporal and cell type-specificity and instead attempt to broadly restrict excitability that can at best mask symptoms while resulting in various side-effects^5^. Recently, a number of experimental studies demonstrated that it is possible to control spontaneous chronic seizures and comorbidities through closed-loop interventions. Importantly, these studies targeted specific cell ensembles in given brain regions and delivered intervention stimuli selectively only at particular times, all with minimal side-effects^6–9^. Although these experiments targeted groups of cells, the most desirable target for intervention would be single cells that exert maximal control over epileptic networks. In this study, we searched for a novel class of potential neuronal targets that could serve as maximally selective controllers for interventions to stabilize epileptic dynamics. To achieve this ambitious goal, we sought to uncover new features of cell-cell communication networks extracted from experimental data that are essential for pathological seizure dynamics to emerge.

One conserved feature of complex networks is the presence of richly connected yet sparse hub neurons^10^. It has been shown that these neurons are critical for influencing network dynamics in biological neural circuits^11–13^. Specifically, experimental studies have shown that hub neurons orchestrate network synchrony in the developing brain^12^ and maintain their effectiveness in adulthood^13^. Simulations in large-scale data-driven hippocampal dentate gyrus computational model of temporal lobe epilepsy (TLE) predicted that perturbation of a population of neurons that included hubs was sufficient to initiate a seizure^11^. These efforts have sparked significant interest in targeting hub neurons for effective seizure control^7,9^. While these hub neurons are traditionally defined based on connection count (i.e., they are unusually richly connected to other cells), there is current interest in elucidating similarly rich higher-order connectivity patterns called motifs that are present in complex networks and are widely believed to inform network function^14,15^. Therefore, it remains an open question whether improved targets for selective seizure control can be identified based on the higher-order network structure of epileptic circuits.

To address the latter question, we deployed whole-brain cellular resolution calcium imaging to capture neural dynamics in larval zebrafish, whose neuronal circuitry shares many conserved features with mammals^16^ and thus have been instrumental in basic^17,18^ and translational^18,19^ epilepsy research. Imaging was performed in a well-characterized zebrafish model of acute seizures^20^. The underlying cell-cell effective connectivity (i.e. communication) networks for baseline and preseizure neural dynamics were extracted^21,22^ and biologically constrained to the zebrafish neuroanatomical connectome for the first time^23^. Simulated perturbation of a single traditional hub neuron significantly destabilized preseizure networks compared to baseline networks. Higher-order analysis^24^ on traditional hubs revealed that network instability in epileptic circuits is causally linked to a subset of hubs whose surrounding neighborhood is rich in feedforward motifs, enhancing downstream excitation. Disconnecting such superhub neurons robustly stabilized networks to perturbation, even though superhubs did not have the highest connection count among the broader hub cell class.

Importantly, similar results were also found in the hippocampal dentate gyrus of chronically epileptic mice compared to control, indicating that our key findings hold in the mammalian brain and in chronic temporal lobe epilepsy (TLE), the most common epilepsy in adults. Collectively, these results identify the emergence of superhubs as a critical step in the destabilization of epileptic circuits, advancing the goal of achieving maximally selective single-cell control of epilepsy with minimal side effects. Looking into the future, studying network interactions at cellular scale opens avenues of investigation that seek to unify single-cell dynamics with system-level communication in both control and pathological brains.

## Results

### Whole brain imaging of larval zebrafish acute seizure model at single cell resolution

Whole brain larval zebrafish (*Tg(elavl3*:*H2B-GCaMP6s);* N=3) imaging was performed with volumetric two-photon microscopy for 25 minutes at 2Hz capture rate (**Fig 1A**, see Methods). Five minutes of baseline data was recorded for each fish followed by 15 mM PTZ^17,20^ bath application, which blocks inhibitory GABAA conductance. Each fish exhibited at least one seizure prior to cessation of imaging. LFP recording was not included as previous studies have shown robust correlation between calcium signal and field potentials in PTZ model^17^. Anatomical volume stacks were registered and neural somata (**Fig 1B**) were extracted with methods used in^16^. We extracted fluorescence time series from 5000-7000 active neurons per fish across all major brain regions (**Fig 1C**).

**Figure 1:**
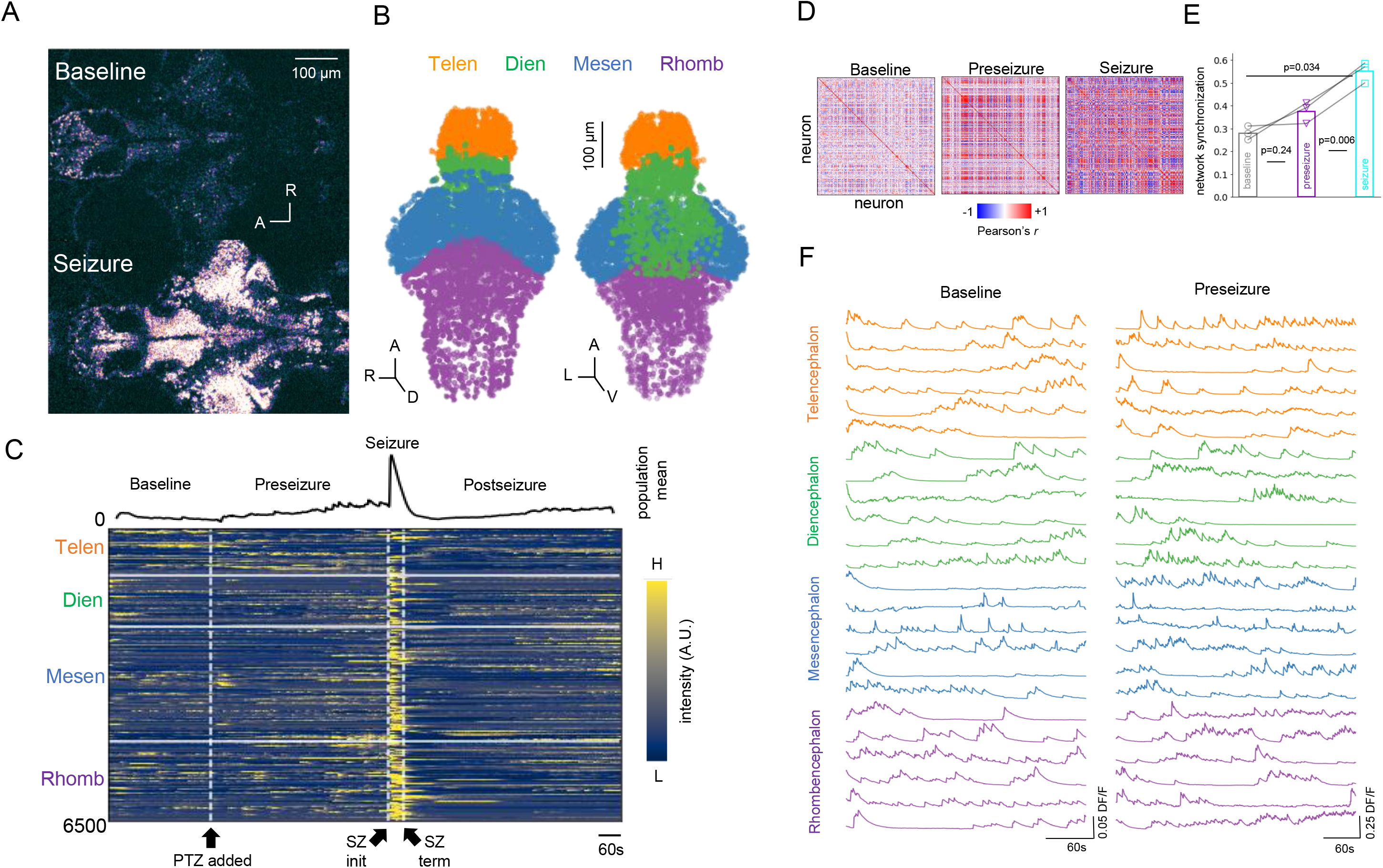
Whole brain imaging of larval zebrafish acute seizure model at single-cell resolution. **A**: Representative z-plane images acquired from whole-brain cellular resolution two-photon microscopy of larval zebrafish before PTZ (baseline; top) application and during PTZ-induced seizure (bottom). **B**: Extracted neural somata point cloud. Colors indicate major brain regions. Orange: telencephalon; Green: diencephalon; Blue: mesencephalon; purple: rhombencephalon **C**: (top) Population mean calcium signal and (bottom) heatmap of single-cell functional calcium dynamics from neurons extracted in B. PTZ application, seizure initiation, and seizure termination are demarcated by arrows. Imaging was performed for 25 minutes and PTZ was added 5 minutes into the imaging session. **D:** Cross-correlation matrices of single-cell calcium dynamics during baseline, preseizure, and seizure epochs. **E:** Quantification of network synchrony from correlation matrices in D show that single-cell calcium dynamics during seizure epoch are significantly more synchronized compared to single-cell calcium dynamics in baseline (one-sided paired t-test, p=0.034 after Bonferroni correction) and in preseizure (one-sided paired t-test, p=0.006 after Bonferroni correction) epochs. **F:** Single-cell calcium traces over major anatomical regions plotted during baseline and preseizure epochs. Note the differences in vertical scale bars.

The start of the preseizure state was defined as 1 minute after PTZ is introduced to account for transient activity. Seizure initiation was defined as three standard deviations above population mean calcium signal during baseline (pre-PTZ) and termination was defined as when the population mean signal dropped below this threshold. In agreement with prior literature^17^, calcium dynamics within detected seizure period lasted 35-40 seconds and displayed significantly higher synchronicity (i.e. hypersynchronous state) compared to baseline (one-sided paired t-test, p-adjusted=0.034) and preseizure (one-sided paired t-test, p-adjusted=0.006) epochs (**Figs 1D,E**). For this work, we will be modeling baseline and preseizure calcium dynamics (**Fig 1F**), and we will not be focusing on seizure or post-seizure periods.

### Cellular resolution effective connectivity modeling to extract cell-cell communication networks

Effective connectivity modeling to extract cell-cell communication networks was performed with chaotic recurrent neural networks (RNN)^21^ (**Fig 2A**, see Methods), which has previously been successful in fitting cellular-resolution calcium dynamics in the zebrafish^25^. Each neuron imaged experimentally is represented by a node in the model. The parameters of the model are the synaptic connections between nodes, interpreted as how much causal influence a source has on its target over a sub-second temporal window and can vary in sign and magnitude. The self-perpetuating chaotic dynamics were controlled through FORCE learning^22^ (Methods), which uses a recursive-least square optimization on the connectivity parameters to reproduce a specified target output **(Fig 2C)**.

**Figure 2:**
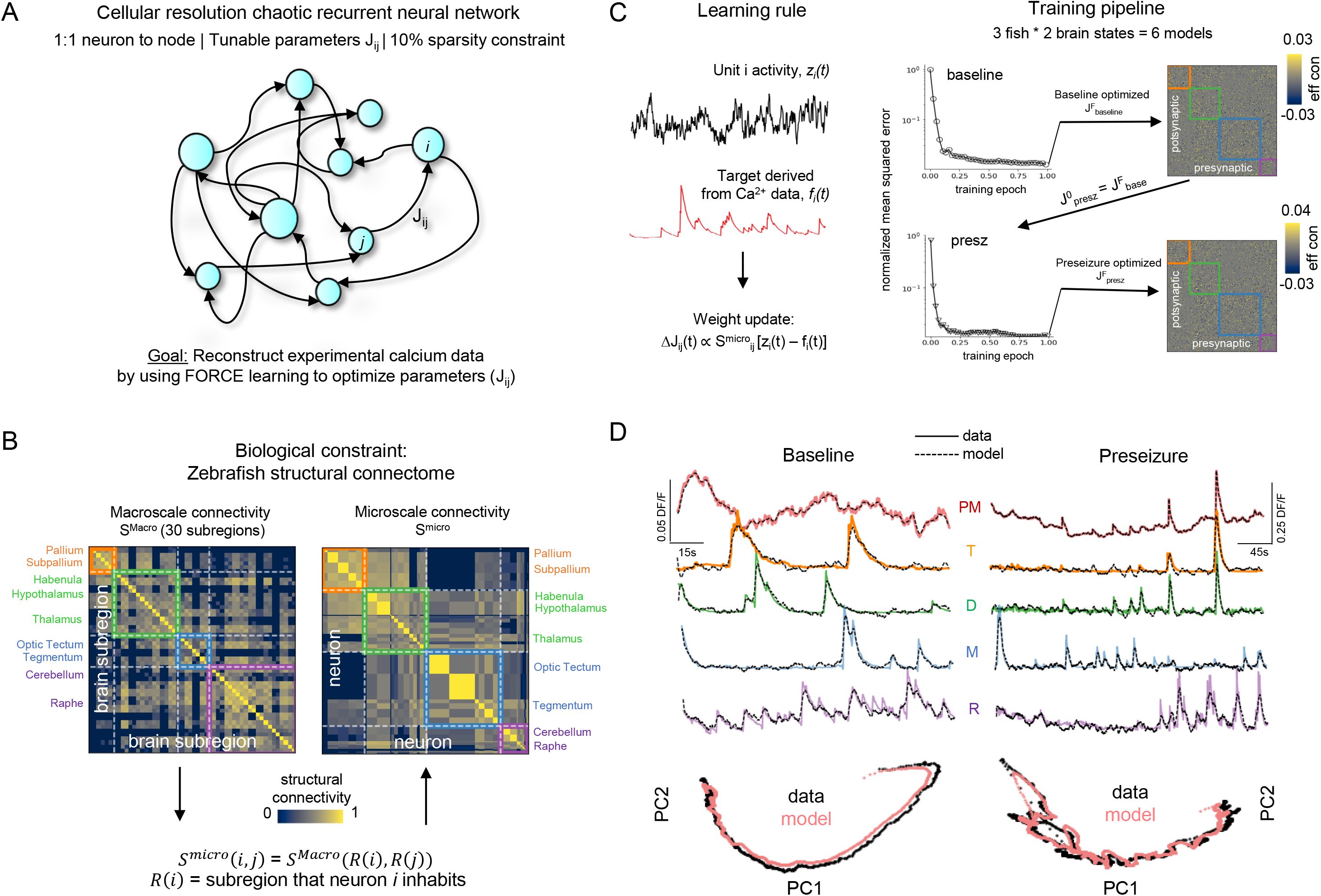
Cellular resolution effective connectivity modeling to extract cell-cell communication networks. **A:** Cellular resolution chaotic recurrent neural network (RNN) with 90% sparsity constraint to prevent overfitting. Each neuron imaged in the larval zebrafish is represented as a node in the RNN. Edges represent parameters of the model and are were optimized with FORCE learning to match experimental calcium data. **B:** The zebrafish structural connectome was incorporated as a biological constraint. (Left) Zebrafish macroscale connectivity matrix (S^Macro^). (Right) Zebrafish microscale connectivity (S^micro^), which represents the strength of connectivity between the regions in which neurons *i* and *j* occupy. **C:** (Learning rule) FORCE learning tunes the weight J_ij_ between neuron i (target) and j (source). The update is proportional the structural connectivity score S^micro^_ij_, multiplied by the difference between unit activity of node *i* (black trace) and target Ca^2+^ waveform acquired experimentally (red trace). (Training pipeline) Models converged using the weight update that incorporated the structural connectome. The baseline model was trained on baseline calcium dynamics with an initial random matrix. To map the changes to the underlying microcircuit connectivity resulting from bath wash in of PTZ, the optimized parameters of the baseline model was then used as the seed for training the preseizure model on preseizure calcium dynamics. **D:** Representative examples of mean population Ca^2+^ trace and individual Ca^2+^ traces with modeled fits overlaid for baseline (left) and preseizure (right) dynamics. Note the scale bars. P: population mean calcium signal. (Bottom) PCA state-space analysis of experimental (black) and modeled calcium activity (red).

Target outputs were the experimentally acquired calcium dynamics. We furthermore employed a 10% sparsity constraint 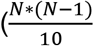 total parameters) to match reported connectivity patterns^26^ and to prevent overfitting. One major issue is whether local minima visited during optimization lead to parameter sets that are realistic given the underlying neuroanatomy. To mitigate this, we constrained the optimization procedure using the known zebrafish structural connectome^23^ (**Fig 2B**, Methods). Specifically, weights are adjusted proportional to how strongly connected the regions in which the source soma and target soma inhabit (**Fig 2C**, **left**). Preseizure networks were optimized from best-fit baseline networks in order to untangle cell-cell effective rewiring caused by PTZ (**Fig 2C**, **right**). For consistency, this was repeated three times with different initial conditions for each fish. Models converged for both baseline and preseizure state (**Fig 2C**, **right**) and FORCE learning captured individual and global calcium dynamics (**Fig 2D**, note scale bars).

Community detection (Methods) on the constrained optimized parameter matrix identified relevant macroscale anatomical structures which were not observed in unconstrained models (**Supplementary Figure 1**). Furthermore, model dynamics were primarily driven by synaptic transmission and not noise (**Supplementary Figures 2A,B**). Outgoing inhibition per neuron decreased after PTZ bath wash-in (**Supplementary Figure 2C**), consistent with PTZ reducing inhibitory conductances^27^.

### Identification of outgoing and incoming traditional hubs

Traditional hub neurons were segregated into two classes based on connection count: incoming and outgoing (**Fig 3A**). Outgoing hubs project while incoming hubs receive numerous strong connections. The optimized parameter matrix was binarized by keeping only the strongest excitatory connections (top 10%; **Fig 3B**). Degree distributions for both hub subtypes and across brain states displayed heavy-tailed distributions as reported experimentally^12^ (**Fig 3C**). To identify outgoing and incoming hubs, the 90^th^ percentile cutoff was calculated (i.e., hubs were defined as the 10% most connected cells; note that changing the 90% cutoff to a different percentile did not alter the key findings, see below). Cutoff values were independently calculated for incoming and outgoing hubs and between baseline and preseizure distributions (**Fig 3C**).

**Figure 3:**
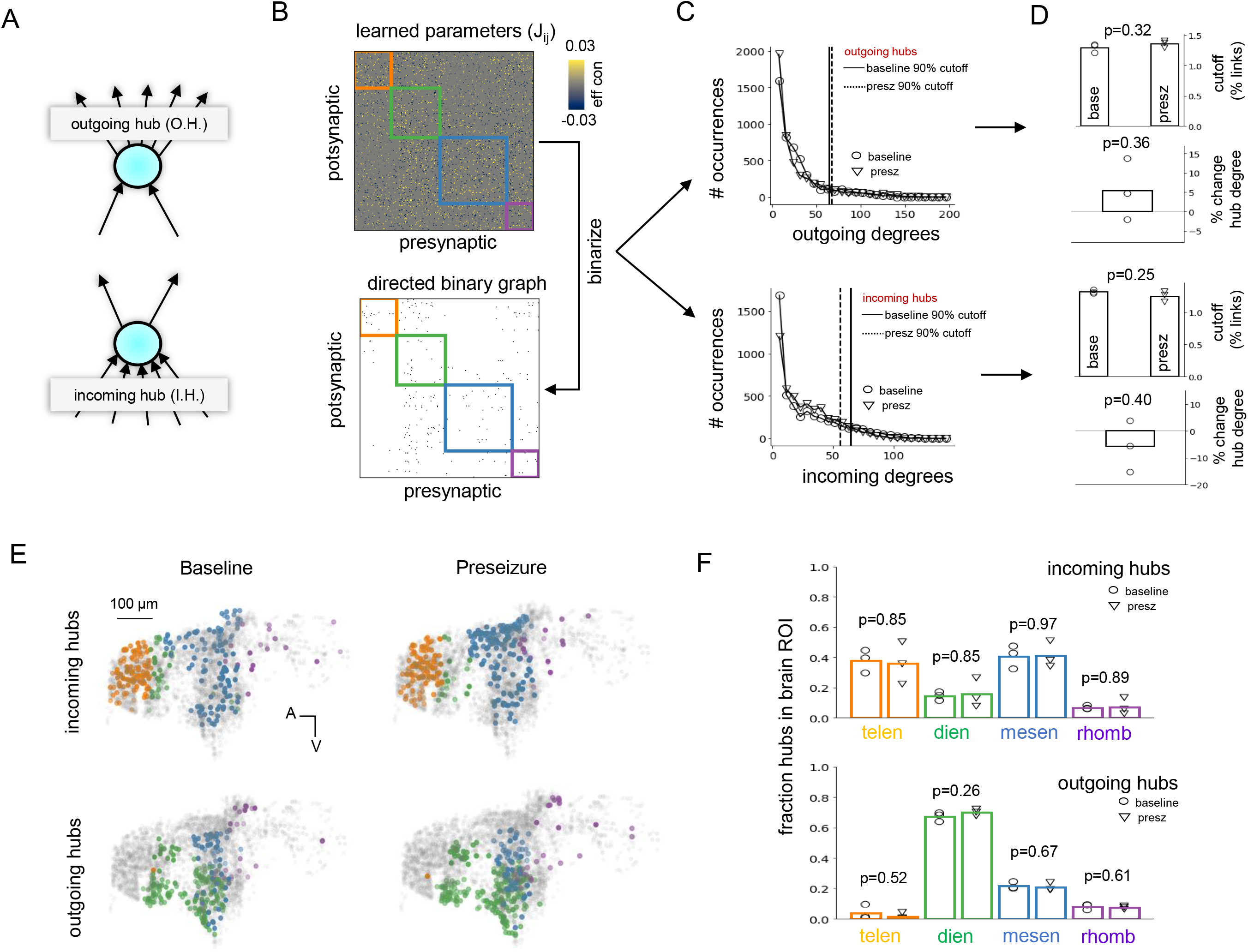
Identification of outgoing and incoming. **A:** Hub neurons were separated into incoming (numerous and strong postsynaptic inputs) and outgoing (numerous and strong presynaptic outputs). **(B,C)**: Algorithm for identifying incoming and outgoing hub neurons in parameter matrix optimized though constrained FORCE learning. **B:** The parameters (top) are binarized into a 0-1 graph (bottom) by keeping the top 10% of excitatory weights. Then, the incoming and outgoing degree for each neuron is calculated from the binarized graph. **C:** Baseline and preseizure network degree distributions for outgoing (top) and incoming (bottom) degree are heavy-tailed, resembling a power-law. A 90% cutoff (vertical lines) was used to identify outgoing and incoming hubs in each network. **D:** 90% threshold cutoff as a fraction of nodes in network and percent change of average hub degree as measured from outgoing (top) and incoming (bottom) degree distributions. Neither paramters was statistically significant between baseline and preseizure networks (two-sided unpaired t-test, p>0.05). **E:** Spatial distribution of incoming and outgoing hubs for baseline and preseizure states. Orange = telencephalon; Green = diencephalon; Blue = mesencephalon; Purple = rhombencephalon. **F:** Fraction of incoming (top) and outgoing (bottom) hubs residing in each macroscale brain region for baseline (open circle) and preseizure (open triangle) networks. Incoming hubs were consistently localized to telencephalon and mesencephalon. Outgoing hubs were consistently localized to diencephalon. Baseline and preseizure networks had similar macroscale spatial organization of incoming and outgoing hubs (unpaired t-test, p>0.05).

Importantly, outgoing and incoming hub cutoff threshold, and mean outgoing (incoming) degree of outgoing (incoming) hubs were not statistically different (two-sided unpaired t-test, p>0.05) between baseline and preseizure states (**Fig 3D**). There was approximately a 5% difference in the average outgoing (incoming) degree of outgoing (incoming) hubs between baseline and preseizure networks (**Fig 3D**). The spatial locations of outgoing and incoming hub neurons were visualized in the zebrafish anatomy for both baseline and preseizure networks (**Fig 3E**). Outgoing hubs were primarily located in diencephalon and incoming hubs were primarily located in mesencephalon and telencephalon **(Fig 3F)**. PTZ did not cause significant changes in the macroscale anatomical locations of outgoing and incoming hubs **(Fig 3F)** (two-sided unpaired t-test, p>0.05).

### Perturbation of individual outgoing hubs destabilize preseizure networks

Chaotic recurrent neural networks are a generative model and can therefore create synthetic calcium traces for each cell. Single-cell perturbation simulation studies are of growing importance for studying functional properties of large neural populations and have to date elucidated how activity is coordinated in recurrent cortical networks^28^. With this in mind, we tested the hypothesis that preseizure networks are more sensitive to perturbation of a single hub neuron compared to baseline networks. Simulated perturbations involved 500 ms depolarizing current injection into a single hub neuron after 20% of epoch duration had elapsed. Depolarization of an outgoing hub increased activity in neurons receiving strong excitatory inputs from the perturbed neuron and this increase was higher in preseizure state than baseline state (**Supplementary Figures 3A,B**). Considering this, we quantified the effect of perturbation on global network dynamics by measuring the ‘trajectory deviation’ (time-normalized Euclidean distance, see Methods) between the mean population calcium signal of the unperturbed network with the mean population calcium signal of the perturbed network.

Depolarization a single outgoing hub in the preseizure state caused significantly higher deviation (one-sided Mann-Whitney U-test. p<0.001) in global dynamics (**Fig 4B,C**) compared to equivalent simulations in baseline networks (**Figs 4A,C**). Depolarizing incoming hubs (**Supplementary Figures 3C,D, Fig 4C**) and non-hubs (**Supplementary Figures 3C,D**) had significantly less influence in both networks. Incoming hub perturbation affected dynamics more greatly (one-sided Mann-Whitney U-test, p<0.001) in preseizure networks (**Fig 4C**). Therefore, we normalized the outgoing hub population by the median trajectory deviation of the incoming hub population. The data maintains that perturbation of a single outgoing hub in the preseizure state altered global dynamics more significantly within individuals (one-sided Mann-Whitney U-test, p<0.001) (**Fig 4D**) and across the population (one-sided paired t-test, p=0.041) (**Fig 4E**). Taken together, these data suggest that preseizure networks are more sensitive to perturbations and that even a single outgoing hub neuron can significantly influence global dynamics.

**Figure 4:**
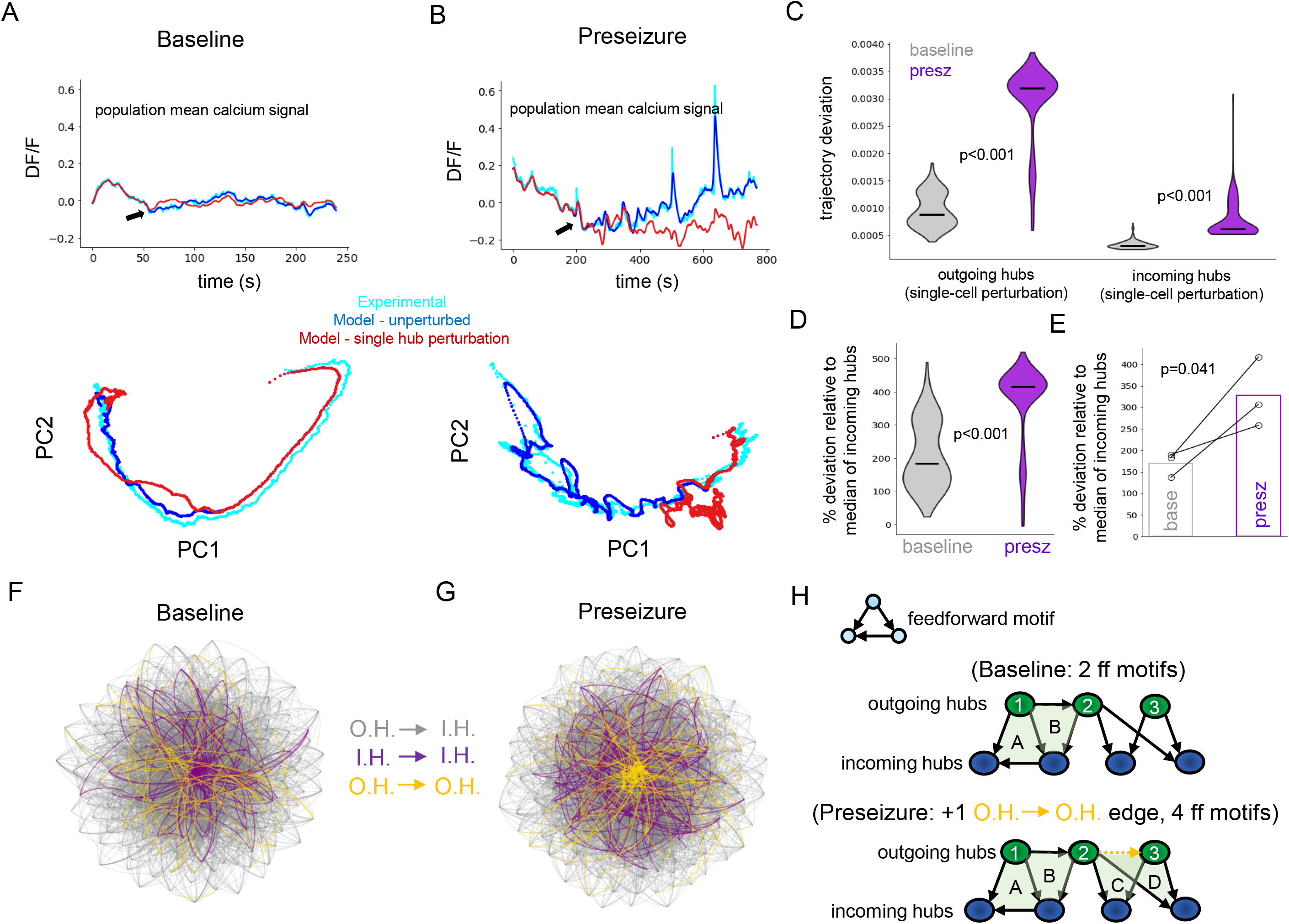
Perturbation of individual outgoing hubs destabilize preseizure networks. **A:** Control network response to perturbation (black arrows) of a single outgoing hub. (Top) Population mean calcium signal of experimental data (cyan), unperturbed model (blue), and perturbed model simulation (red). (Bottom) PCA analysis reveals little change in network dynamics in response to perturbation. **B:** Perturbation of single outgoing hub in preseizure network showing significant changes to network dynamics compared to A. **C:** Violin plots show that perturbing individual outgoing hubs (one-sided Mann-Whitney U-test, p<0.001) and incoming hubs (one-sided Mann-Whitney U-test, p<0.001) in preseizure networks (purple) had significantly higher influence on network dynamics compared to similar simulations in baseline networks (gray). **D:** Outgoing hub trajectory deviation distributions in baseline (gray) and preseizure networks (purple) normalized by incoming hub median trajectory deviation score for the respective populations. Perturbation of outgoing hubs in preseizure state has significantly more influence over network dynamics (one-sided Mann-Whitney U-test, p<0.001). **E:** Median values from (D) were extracted for each fish and plotted, revealing that preseizure networks have significantly reduced resiliency to perturbation of a single outgoing hub (one-sided paired t-test, p=0.042). **F,G:** Visual representation of connections between outgoing and incoming hubs for baseline (F) and preseizure (G) network after constrained FORCE learning. Graphs were generated using Barnes-Hut algorithm^53^. Gray edges: outgoing hubs (O.H.) to incoming hubs (I.H.); Purple edges: incoming hubs (I.H.) to incoming hubs (I.H.); Golden edges: outgoing hubs (O.H.) to outgoing hubs (O.H.). **H**: Toy model of connections between outgoing (green) and incoming (blue) hubs for baseline (top) and preseizure (bottom) networks.

### Visualizing connections between outgoing and incoming hubs

PTZ bath wash-in did not significantly change several parameters associated with the outgoing and incoming degree distributions (**Figs 3C,D**) and did not significantly reorganize which macroscale anatomical regions the hub cells occupied (**Figs 3E,F**). Therefore, we hypothesized that nuanced rewiring of microcircuit effective connectivity patterns may be critical for understanding increase sensitivity to perturbation (**Fig 4**). We visualized the connectivity patterns between outgoing and incoming hubs for baseline (**Fig 4F**) and preseizure (**Fig 4G**) networks using force-directed graphs (Methods). Importantly, nodes clustered in force-directed graphs are highly connected. We observed increased number of outgoing – outgoing hub connections (**Fig 4G**, note golden edges forming a ‘core’) in preseizure networks compared to pre-PTZ baseline control.

Enhanced recurrent connectivity between outgoing hub neurons in preseizure networks suggest an important role for motifs. Motifs are subgraphs that are thought to provide insight on the functional properties of complex systems^14,15^. In baseline toy model (**Fig 4H**, **top**), there is a single connection between outgoing hubs and two feedforward motifs present. Based on observations in **Fig 4G**, there are more connections between outgoing hubs. Therefore, preseizure toy model (**Fig 4H**, **bottom**) includes an additional connection between outgoing hubs 2 and 3. As a result of adding a single connection, there are now four feedforward motifs present in the network. Therefore, we hypothesized that the multiplicative emergence of feedforward motifs may be a core feature of cell-cell communication networks that destabilize the preseizure brain.

### Emergence of superhubs in the preseizure brain

Graph clustering is an intuitive method to isolate groups of nodes in a network that form numerous intra-group connections. Specifically, we predict the emergence of superhubs - hubs with interesting higher-order connectivity patterns - using a higher-order local clustering concept. The inputs to the algorithm were the directed binarized graph (**Fig 3B**), a motif *M*, and an individual outgoing hub identified in **Fig 3C** (i.e. the seed). The algorithm finds an optimal cluster surrounding the seed that is rich in *M* with run-time that is invariant of the graph size (**Fig 5A**, **left**). Traditional edge clustering, which only considers simple edges, was also performed as a control (**Fig 5A**, **right**). The edge (motif) conductance metric was used to capture how much information or activity propagates from the traditional (higher-order) cluster to the rest of the network (**Fig 5B**). Optimal edge and higher-order clusters (i.e. minimal conductance) were identified for outgoing hubs in baseline and preseizure networks using the Motif-based Approximate Personalized PageRank (MAPPR) algorithm^24^.

**Figure 5:**
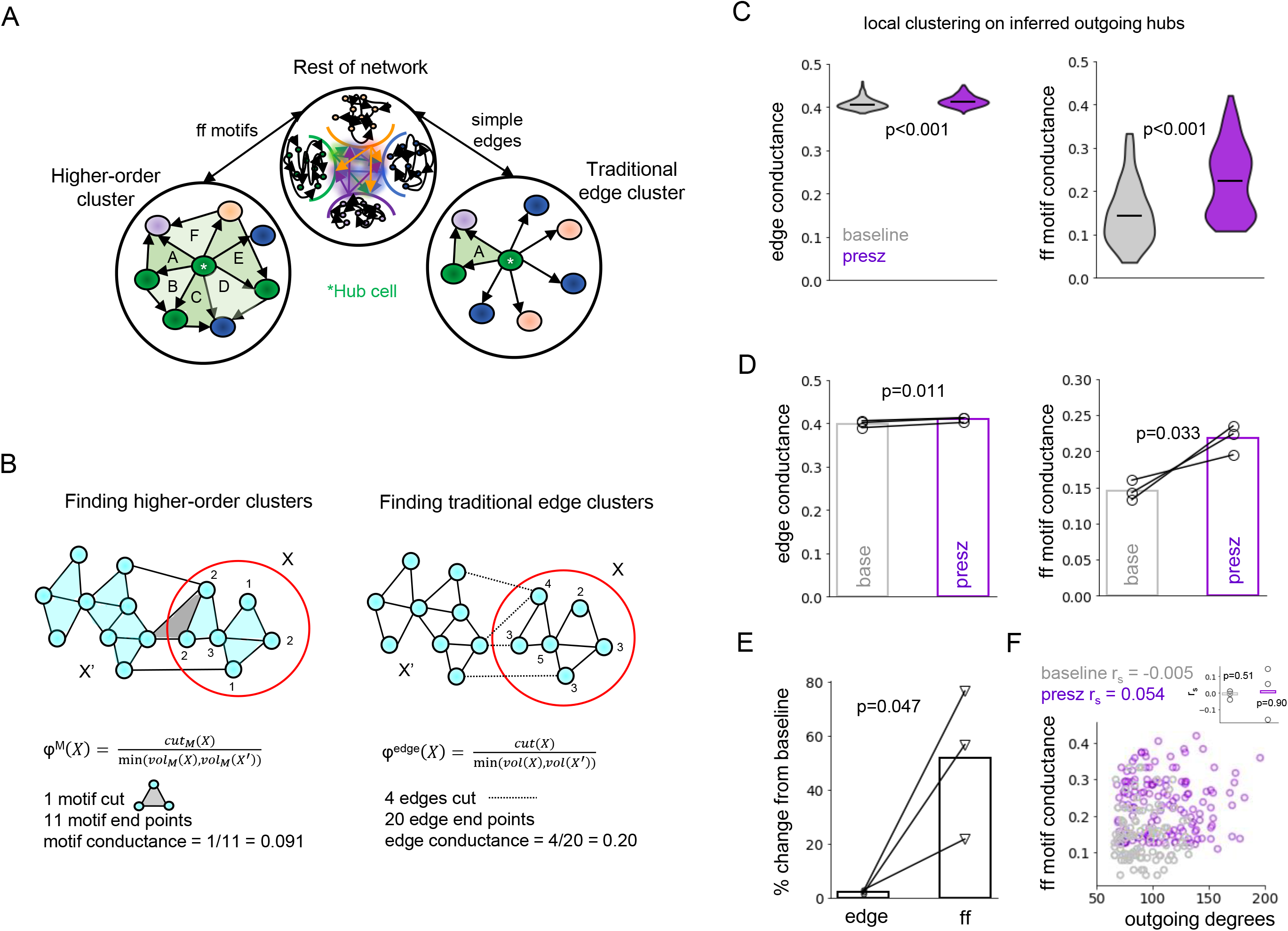
Emergence of superhubs in preseizure brain. **A:** Higher-order clustering enables identification of a collection of nodes that form rich feedforward connections with a single hub neuron (left). This approach contrasts to traditional edge clustering, which only considers simple edges (right). Note that different clusters emerge depending on which method is used. **B:** Toy model of higher-order clustering (left) and edge clustering (right) (from Yin *et al*, 2017). Edge conductance quantifies the cluster quality by considering the ratio of edges that span between partitions and which reside completely inside the partition. The motif conductance metric measures the same ratio but with respect to motifs. Intuitively, the higher the conductance, the more easily it is for excitatory activity to propagate downstream. **C:** Violin plots of edge conductance (left) and feedforward motif conductance (right) of outgoing hub neurons. Both edge (one-sided Mann-Whitney U, p<0.001) and feedforward motif conductance (one-sided Mann-Whitney U, p<0.001) are higher in preseizure than baseline networks. However, note the differences in range. This predicts the emergence of ‘superhubs’ in preseizure networks. **D:** Edge conductance (one-sided paired t-test, p=0.011) and feedforward motif conductance (one-sided paired t-test; p=0.033) are significantly increased in preseizure networks across the sample population. Medians from **C** plotted. **E:** Percent change of feedforward motif conductance relative to baseline was significantly greater than edge conductance (one-sided paired t-test, p=0.047). **F:** Scatter plot of feedforward motif conductance versus outgoing degree of hubs in baseline and preseizure networks reported no significant correlation between the two variables individually or as a group (see inset).

Edge conductance (one-sided paired t-test, p=0.011) and feedforward motif conductance (one-sided paired t-test, p=0.033) were elevated in preseizure networks (**Figs 5C,D**). However, the increase in feedforward motif conductance relative to baseline was significantly higher (one-sided paired t-test, p=0.047) than edge conductance (51.8% vs 2.4%; **Fig 5E**). We performed higher-order clustering on additional motifs, such as cycles, but found very few instances. Therefore, this data suggests that a subset of hubs in preseizure networks allow excitation to propagate more readily to the rest of the network, predicting the emergence of ‘superhubs’. Importantly, both individually (baseline Spearman’s rho=-0.005, p>0.05; preseizure Spearman’s rho=0.054, p>0.05) and as a group (unpaired two-sided t-test, p>0.05), feedforward motif conductance was not correlated to outgoing degree of the outgoing hub (**Fig 5F**).

### Disconnecting superhubs stabilizes preseizure networks

Our evidence so far suggests a correlation between two findings: 1) Perturbation of a single outgoing hub in preseizure networks has significant influence on global network dynamics; 2) A subset of hubs in preseizure networks may be propagate excitatory activity to the rest of the network more easily (i.e. superhubs). To establish a causal link, we performed computational experiments. These simulations involve targeted attacks on hub neurons. The motivation for this is that complex networks in nature are vulnerable to attacks on hub neurons^29^, and which has been validated in neural circuits through closed-loop optogenetic studies^6,7^.

We first compared the role of feedforward motifs versus simple edges on network function. Edges that projected from outgoing hubs and targeted neurons in its local higher-order cluster or its local edge cluster were damped by a factor ranging from 0 (disconnect) to 1 (no change) (**Fig 6A**). Dampening the outputs of outgoing hubs to their respective higher-order cluster has greater effect on network activity in both baseline and preseizure networks (**Fig 6B**, **left**). However, this effect was more pronounced for preseizure state (**Fig 6B**, **right**).

**Figure 6:**
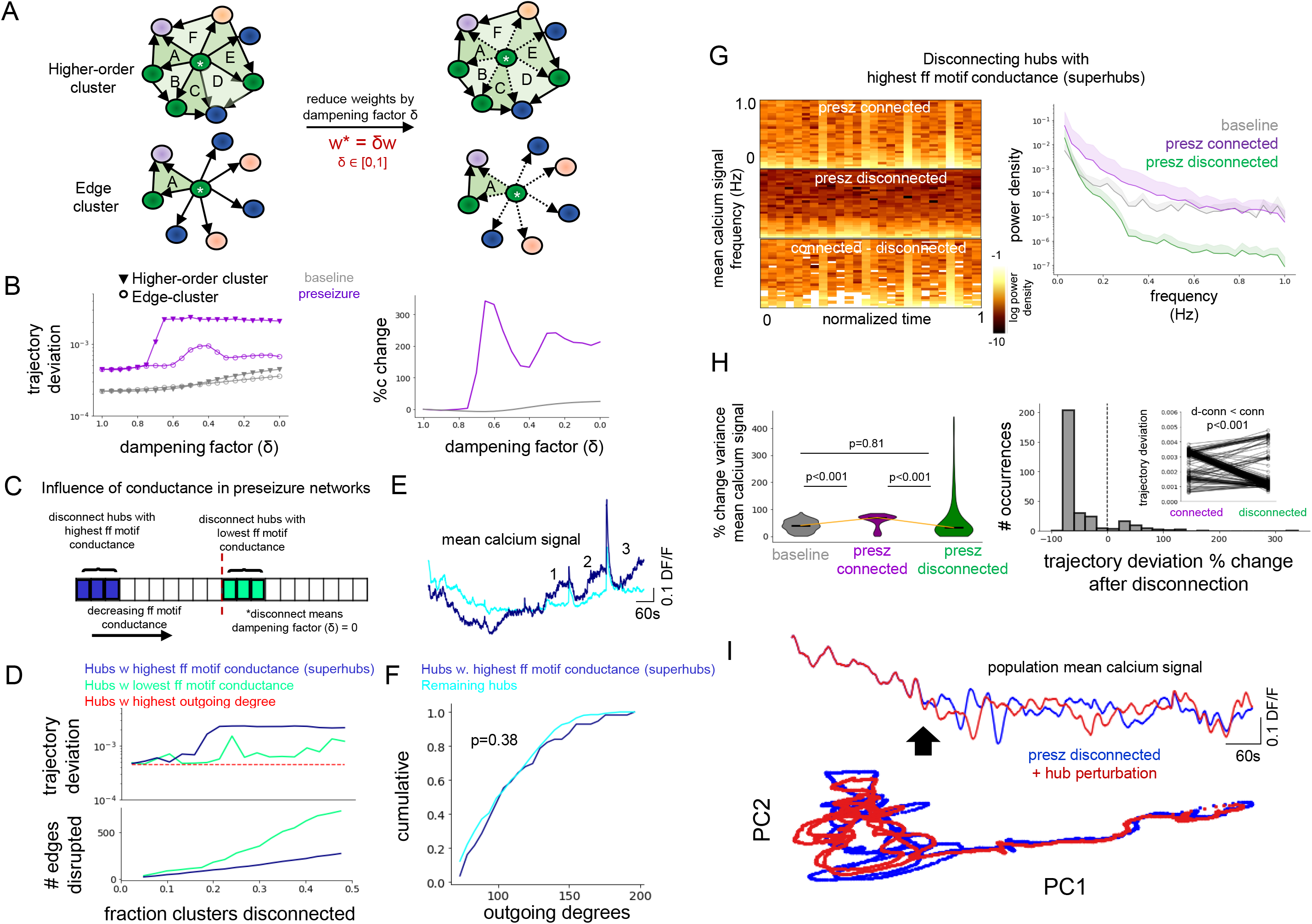
Disconnecting superhubs stabilizes preseizure networks. **A:** Edge weights of all hubs targeting its edge cluster constituents and higher-order cluster constituents were dampened to explore relative importance of simple edges versus edges belonging to feedforward motifs on network dynamics. **B:** (Left) Trajectory deviation after dampening edges. (Right) Trajectory deviation percent change relative to dampening edges from an outgoing hub to its local edge cluster. **C:** Schematic for testing the effect of disconnecting hubs with the highest conductance versus lowest conductance on preseizure network dynamics. **D:** (Top) Trajectory deviation versus fraction of clusters disconnected in the highest conductance and lowest conductance partitions. (Bottom) Number of edges disconnected for each group. **E:** Mean calcium signal of high conductance higher-order hubs (i.e. superhubs) and the remaining hub population. **F:** Outgoing degree cumulative distribution for superhubs versus remaining hub population. Importantly, superhubs are not biased towards the highest outgoing degrees. **G:** (Left) Mean population calcium signal spectrograms generated from unchanged/connected preseizure network (top), preseizure network with superhubs disconnected (middle), and difference. (Right) Power density spectrum (PSD) of mean population calcium signal for baseline, connected preseizure (violet), and disconnected preseizure network. **H:** (Left) Percent change of signal variance (i.e. total power) in mean population calcium signal measured before and after perturbation of a single hub. (Right) Trajectory deviation percent change. **I:** (Top) Mean population calcium signal of disconnected preseizure network before and after perturbation of a single outgoing hub. (Bottom) PCA analysis before and after perturbation.

Second, we validated that hubs with the highest motif-conductance (i.e. could propagate activity downrange more easily) have greater influence over network dynamics than hubs with low network conductance. Hubs in preseizure network were ranked based on their feedforward motif conductance score and split into either top half or bottom half. Within each half, an equivalent fraction of higher-order clusters was disconnected from the network (**Fig 6C**; dampening factor = 0.0). Simulations confirmed that disconnecting hubs with the highest conductance was effective in changing global network activity (**Fig 6D**, **top**). Interestingly, this effect was observed despite more edges being removed in the bottom half group (**Fig 6D**, **bottom**), and in total represented less than 0.1% of all edges in the network. Brute force disconnecting hubs with the highest outgoing degree did not alter global network dynamics (**Fig 6D**, **top**). Hubs with highest conductance displayed increasing activity in between high calcium events compared to the remaining higher-order hub population (**Fig 6E**, note numbers) and were not biased towards having the highest outgoing degree values (one-sided KS test. p>0.05) (**Fig 6F**). Therefore, motif conductance gives important insight on network function and that this insight could not have been revealed through mining the network for simple edges. These hubs with the highest motif conductance are now explicitly labeled as ‘superhubs’.

Third, we tested the hypothesis that superhubs rendered the network more stable to perturbation. Superhubs - hubs with the highest 37.5% feedforward motif conductance scores - were disconnected (**Fig 6D**, **top**). Oscillatory power was dampened uniformly over all frequency bands for most measured time periods (**Fig 6G**). As expected, the percent change in signal variance (i.e. total power) in response to perturbation of a single outgoing hub in the fully connected preseizure network was significantly higher than baseline network (39.8% to 69.1%; one-sided Mann-Whitney U-test, p-adjusted<0.001). Disconnecting superhubs from the preseizure network significantly reduced percent change in signal variance from 69.1% to 31.7% (one-sided Wilcoxon signed-rank test, p-adjusted<0.001) (**Fig 6H**, **left**). Furthermore, percent change in signal variance of the disconnected network was not significantly different compared to baseline network (two-sided Mann-Whitney U-test, p-adjusted=0.81) (**Fig 6H**, **left**), evidence of renormalization. Disconnecting superhubs significantly decreased the trajectory deviation of global network activity compared to fully connected preseizure network (one-sided Wilcoxon signed-rank test, p<0.001) (**Figs 6H,I**). As a control (**Supplementary Figure 4**), disconnecting superhubs reduced the percent change in signal variance as a response to perturbation more significantly than either disconnecting the same number of hubs randomly (31.7% versus 99.8%, one-sided Wilcoxon signed-rank test, p<0.001) or the same number of low-conductance hubs (31.7% versus 67.6%; one-sided Wilcoxon signed-rank test, p<0.001).

Therefore, the network is more resilient to perturbation when disconnecting superhubs. Taken together, these three computational experiments reveal that the microcircuit connectivity architecture surrounding hub neurons is a critical novel feature of stability in epileptic networks.

### Superhubs in a mouse model of chronic temporal lobe epilepsy

Converging evidence has found similar statistical properties in pathological single-cell dynamics from larval zebrafish and mouse hippocampal dentate gyrus^17,30,31^. We therefore performed the same analysis in healthy control (N=3) and epileptic mice (intrahippocampal KA model; N=3) to address the question of whether our key findings in zebrafish acute seizure model held in the mammalian brain circuit and in a chronic model of TLE, the most prevalent form of epilepsy in adults. Mice expressed GCaMP6f in dorsal dentate gyrus (DG) and the epileptic group had kainic acid (KA) injected unilaterally into ipsilateral ventral hippocampus. 2p calcium imaging of DG granule cells was performed in healthy mice and in chronically epileptic mice (**Figs 7A,B**, note differences in vertical axis of scale bars).

**Figure 7:**
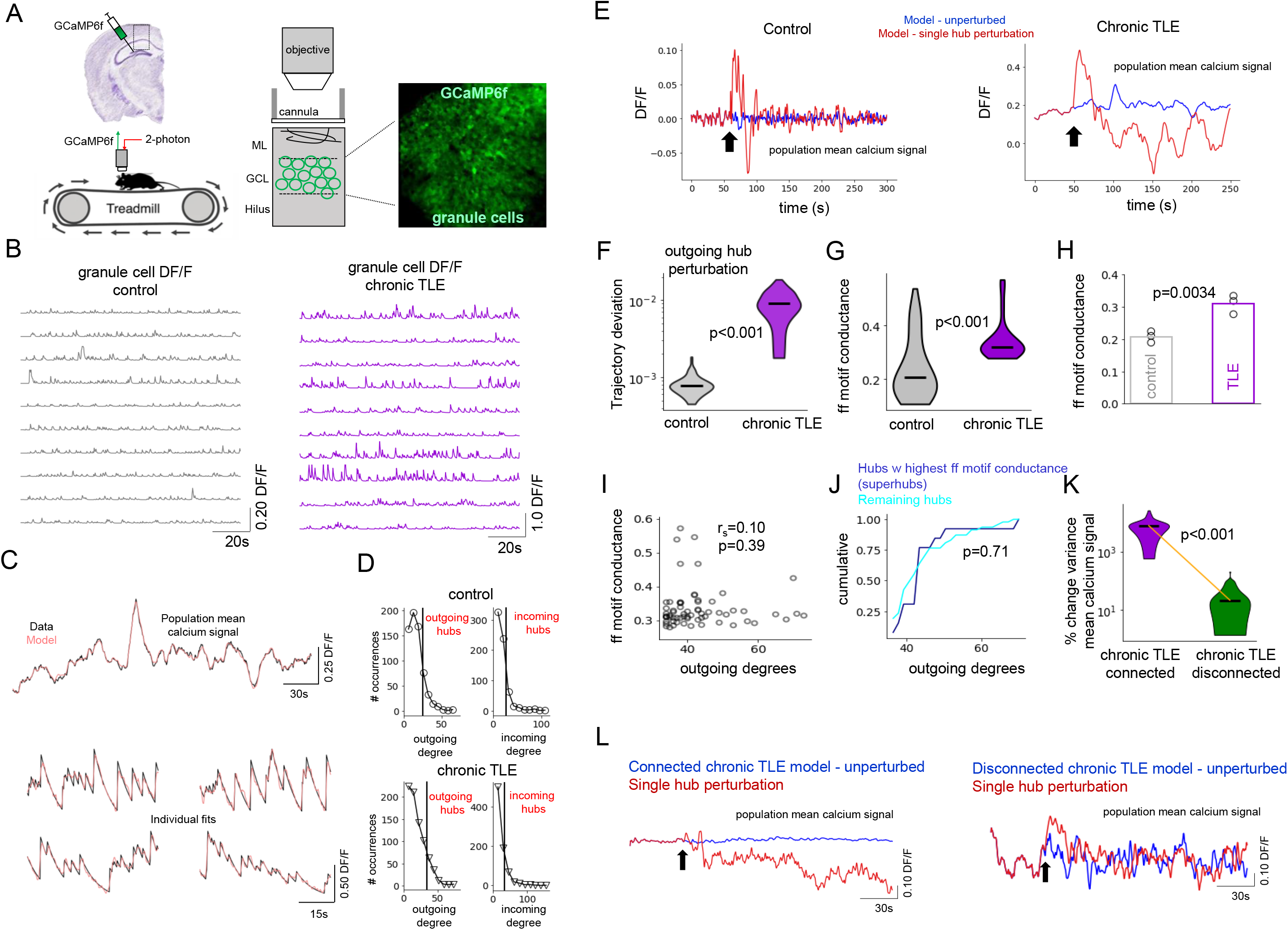
Superhubs in kainic acid mouse model of chronical temporal lobe epilepsy. **A:** Experimental setup. Control and chronically epileptic mice were virally injected with GCaMP in dentate gyrus (DG) and imaged with 2p microscope. **B:** DG granule cell DF/F for control and chronically epileptic mouse. **C:** Models fits of experimentally recorded granule cells from chronically epileptic granule mouse using FORCE learning. **D:** Outgoing and incoming degree distributions for modeled control dentate and chronically epileptic dentate networks showing heavy-tailed distributions. **E:** Network response to perturbation (black arrows) of a single outgoing hub neuron in a modeled control (left) and chronically epileptic (right) network. **F:** Trajectory deviation in response to perturbation of individual outgoing hubs is significantly higher (one-sided Mann-Whitney U-test, p<0.001) in control (gray) than chronically epileptic dentate network (purple). **G:** Feedforward motif conductance of individual hubs is significantly higher (one-sided Mann-Whitney U-test, p<0.001) in control (gray) than chronically epileptic (purple) dentate network. **H:** Feedforward motif conductance is significantly increased in chronically epileptic networks across the sample population (one-sided unpaired t-test, p=0.0034). **I:** Outgoing degrees and feedforward motif conductance are not significantly correlated. **J:** Hubs with highest feedforward motif conductance values (i.e. superhubs) did not have the highest outgoing degrees (one-sided KS-test, p=0.71). **K:** Disconnecting feedforward edges of superhubs significantly reduced the response of a chronically epileptic dentate network to perturbation, as measured by percent change in global signal variance (one-sided Wilcoxon signed-rank test, p<0.001). **L:** Population mean calcium dynamics before (blue) and after (red) perturbation of a single hub for fully connected (left) and disconnected (right) chronically epileptic dentate network.

Granule cell calcium dynamics were modeled in for both groups of mice (600-800 neurons/mouse; 3-5 minute window; **Fig 7C**) and incoming and outgoing hubs were identified (**Fig 7D**) with similar methods used in zebrafish. Modeled control networks were more resilient to single outgoing hub perturbation than chronically epileptic dentate network (**Figs 7E,F**). Global network dynamics were significantly less affected (one-sided Mann-Whitney U-test, p<0.001) when perturbing incoming hubs compared to outgoing hubs for both networks (**Supplementary Figures 5A,B**).

Similar to zebrafish, feedforward motif conductance of local higher-order clusters seeded on outgoing hubs was significantly higher (one-sided unpaired t-test, p=0.0034) in chronically epileptic dentate networks compared to control dentate networks (**Figs 7G,H**). Feedforward motif conductance was not correlated to outgoing degree (Spearman’s rho=0.10, p>0.05) (**Fig 7I**) and top high conductance hubs were not biased towards the highest outgoing degree values (one-sided KS test, p>0.05) (**Fig 7J**). We disconnected superhubs - hubs with the top 20% highest feedforward conductance - in chronically epileptic dentate networks. Single hub perturbation simulations showed significant reduction (one-sided Wilcoxon signed-rank test, p<0.001; **Supplementary Figure 5C**) in global network trajectory deviation and in change in global network signal variance (one-sided Wilcoxon signed-rank test, p<0.001; **Fig 7K.** This reduction was more significant compared to disconnecting the same number of random hubs (one-sided Wilcoxon signed-rank test, p=0.0077) or low-conductance hubs (one-sided Wilcoxon signed-rank test, p=0.0024) (**Supplementary Figures 5D,E**). Example traces of global network dynamics from TLE network in response to perturbation of the same hub before and after disconnecting superhubs are presented in **Fig 7L**.

These similar findings across both acute seizure model in zebrafish and in chronically epileptic mice suggest that the emergence of superhubs is a core principle of network reorganization that may be a key feature contributing to the destabilization of the epileptic brain. Targeting these superhubs may be more effective in cellular scale control of epileptic circuits then hubs defined traditionally on connection count.

## Discussion

Here, we sought to characterize the connectivity patterns of cell-cell communication networks in epileptic circuits. Our findings reveal that superhubs, defined through analysis of the higher-order structure of modeled networks extracted from functional calcium imaging data, emerge as key cellular controllers that can powerfully destabilize epileptic circuits. Disconnecting these superhubs was effective in stabilizing networks in larval zebrafish, and, importantly, these results also held true in the dentate gyrus of chronically epileptic mice. Therefore, these results predict the existence of fundamentally novel, single cell targets for maximally selective, minimally invasive seizure control in epilepsy.

### Search for new interventional targets guided by large-scale cellular resolution functional imaging

Current clinical assessment and treatment of epilepsy is to a large extent based on analysis of networks constructed from population recordings such as local field potentials (LFP)^4^ or functional magnetic resonance imaging (fMRI)^32^. However, recent studies have shown that single-cell activity can diverge significantly from what population recordings would predict. For example, apparently self-repeating, macroscopically recurrent epileptiform LFP activity (inter-ictal spikes) indicating network synchrony have been shown to emerge from non-recurrent microscopic single-cell activity. In other words, each inter-ictal event is generated by a different combination of participating cells^17,30,31^. This ‘macro-micro’ disconnect^33^ is a significant challenge for the field and indicates that a more nuanced understanding of pathways at the level of microcircuits is needed to treat the underlying disease^34^. Therefore, in an effort to search for novel forms of maximally selective, ideally single cell-level targets for interventions for seizure control, we performed large-scale calcium imaging at cellular scale and used this as our primary data source for the subsequent modeling and analysis.

### Computational pipeline to study superhubs in cell-cell communication networks

Our computational methodology for uncovering the role of superhubs in destabilizing epileptic circuits was performed using three distinct yet integrated modules: effective connectivity modeling, higher-order network analysis, and single-cell perturbation simulations.

Effective connectivity modeling of macroscopic brain interactions has been employed successfully to gain new insights into brain areas responsible for seizure propagation in the larval zebrafish^35^. We employed chaotic recurrent neural networks^21^ and FORCE optimization^22^ techniques, which have been shown to be effective tools for modeling neural dynamics related to hippocampal sequence generation^36^, motor planning^37^, and coping^25^. Using this approach, we present for the first time a biologically constrained whole brain model of larval zebrafish at cellular resolution, where the biological constraint was incorporated by the inclusion of the zebrafish structural connectome^23^ in the weight-update step. As a result, macroscale subnetworks that resembled major anatomical subregions emerged (**Supplementary Figure 1**). These specific partitions agree with converging evidence that suggest a functional^17,38,39^ separation between the various front and mid-/hind-brain structures.

The parameters of the model form a directed weighted graph, which allowed us to quantify chemoconvulsant- or chronic epilepsy-induced changes to the underlying graph structure using novel methods in network science^24^. We first found heavy-tailed degree distributions of strong excitatory connections for both zebrafish and mouse models (**Figs. 3C**,**8D**)^12^, a feature that is common in large complex networks. This enabled us to identify both incoming and outgoing traditional hub neurons based on incoming and outgoing connection counts. To further analyze the connectivity structure surrounding traditional hubs, we deployed novel higher-order clustering techniques^24^ and quantified motif conductance (**Figs 5A,B**). Importantly, the MAPPR algorithm^24^ identified optimal clusters, which allowed us to directly compare motif conductance between baseline and preseizure networks. While we reported a non-significant increase (~5%) in average outgoing degrees in preseizure networks (**Figs 3C,D**), feedforward motif conductance of higher-order clusters surrounding outgoing hubs increased significantly by 50% on average (**Fig 5E**).

Single-cell perturbation simulations are important tools for generating predictions of how individual cells influence network dynamics^28^. Measures were taken in the modeling step to prevent overfitting, such as including a sparsity constraint on the number of model parameters, which allowed us to perform controlled perturbation simulations. We first interrogated network stability in modeled control and pathological circuits. Results revealed that perturbation of just a single outgoing traditional hub neuron can significantly influence epileptic global network dynamics compared to similar computational experiment in baseline networks (**Figs 4, 7**). One reason for this may be due to loss of inhibition in epileptic circuits^27,40,41^ which normally functions to stabilize network activity^28,42,43^. Taken together, these results highlight that preseizure networks are unstable^28,44^. Additionally, we showed that targeted-attack simulations of identified superhubs (i.e. hubs with high feedforward motif conductance) can stabilize networks to perturbation. These causality-establishing simulations (**Figs 6, 7**) predict that superhubs can more readily propagate sustained excitatory activity downstream in epilepsy compared to control, baseline networks.

### Superhubs in the epileptic brain

The primary conceptual advance in this work was brough to light from mining the higher-order network structure of large-scale cellular resolution networks extracted from experimental neural data. Highly connected but rare hub neurons have been of great interest for studies working towards the goal of advancing seizure control with minimal side-effects^7–9^. This has been motivated by experimental work showing that hub neurons orchestrate network synchrony^12,13^ and by computational work predicting their role in the transition from interictal to ictal discharge^11^.

However, hub neurons are traditionally defined by simple connection counts, a definition that does not consider the rich higher-order features that are often found in complex networks^14,15^. As these ‘motifs’ are thought to influence network function in biological neural circuits^15^, we deployed our computational pipeline towards the goal of identifying potential new targets for control of pathological circuits. The key results presented in this paper could not have been uncovered with traditional lower-order graph mining techniques, as the primary metric to extract superhubs - motif conductance - was not correlated to connection count (**Figs 5F, 6F, 7J**). Furthermore, because disconnecting superhubs stabilized networks, these findings suggest that complex networks can be susceptible to targeted attacks of nodes when considering rich higher-order features rather than simple connection count. This shift in perspective – specifically looking at the patterns of connectivity surrounding hubs as opposed to connections from hubs - may be important for how we interpret the underlying microscale dynamics from macroscale recordings^33^, evaluate the efficacy of anti-epileptic drugs (AEDs)^19^, and develop more strategic closed-loop interventions^6–9^.

Motifs are a critical feature of the structural and functional connectomes^15,26,28^, and it has been hypothesized that the brain maximizes the diversity of functional motifs^15^. This diversity may be related to the phenomenon of criticality (sometimes called ‘edge of chaos’), a state marked by scale-invariant neural avalanches which has been reported in zebrafish^45^. Prior research showed chemoconvulsant-induced disruption of excitation/inhibition (“E/I”) balance in cortical neural networks caused deviation away from criticality (one neuron activates one neuron) and into the supercritical regime (one neuron activates more than one neuron)^46^. In a similar manner, feedforward motifs include a ‘mother’ neuron activating two other cells. Therefore, our discovery of superhubs dense with feedforward connections in the preseizure brain networks suggests a potential new biomarker that may be able to capture a system’s divergence from criticality.

Our findings in the fish brain in an acute seizure model were remarkably similar to the results from the hippocampal dentate gyrus of chronically epileptic mice. While zebrafish brains contain generally similar circuits compared to mice^16^, there are also important differences, for example, concerning cortical structures prevalent in mammals that are often sites of epileptic foci. Furthermore, acute seizure models lack persistent alterations to the underlying genes, ion channels, synapses, and morphological properties that are present in patients with epilepsy. We addressed both of these issues by modeling dentate gyrus granule cell dynamics in mice with chronic TLE. Through our computational analysis pipeline, we were able to show that higher-order interactions held in distinct experimental epilepsy models and in organisms far removed in evolutionary time. Taken together, these results analyzing the higher-order interactions of control and epileptic networks predict a new single-cell target – the superhub - for maximally selective, minimally invasive control of epileptic circuits. Pinpointing particular superhub cells *in vivo* from the functional imaging data using the pipeline presented in this paper is computationally demanding and has not yet been achieved. However, future studies should be able to isolate and manipulate individual superhub neurons in real time through closed-loop interventions as there is a one-to-one correspondence between the model nodes and the biological neurons in the network.

## Methods

### EXPERIMENTAL MODELS AND SUBJECT DETAILS

All procedures were approved by the Institutional Animal Care and Use Committee for both Stanford University and Columbia University.

#### Zebrafish acute seizure model

##### Zebrafish

*Tg(elavl3:H2B-GCaMP6s)*^47^ fish (7 dpf) bred on a Nacre or Casper background were used for imaging and registration. Zebrafish were mounted dorsal side up in a thin layer of 2.5% low-melting point agarose (Invitrogen) in the lid of a 3 mm petri dish (Fisher), using a sewing needle to position the fish under a stereomicroscope (Leica M80). Fish were group-housed under a 14:10 light:dark cycle until the day of experiments, and were fed with paramecia (Parameciavap) twice daily from 5-6 days post fertilization onward. All testing occurred during the late morning and afternoon.

##### Experimental timeline and two-photon imaging

Two-photon volumetric imaging was performed using an Olympus FVMPE multiphoton microscope (Olympus Corporation), with a resonant scanner, in either unidirectional or bidirectional scanning mode. We used a 16x objective (0.8 NA; Nikon). Functional brain imaging was performed at 1.2x zoom (1.44 μm/pix) in 15 z-planes (15 μm spacing) at 0.59s/vol (2500 volumes).

Baseline control imaging was performed for 5 minutes followed by 15 mM bath application of pentylenetetrazol (PTZ) to induce spontaneous seizures^20^. Functional imaging continued for an additional 20 minutes. All zebrafish exhibited at least one putative seizure prior to cessation of imaging.

After functional brain imaging, a structural stack was obtained at 1 μm spacing, starting 15 mm above the first z-plane, ending 15 mm below the last z-plane. Images were registered to Z-brain atlas *Tg(elavl3:H2B-RFP)* volume and single-cell functional calcium dynamics were extracted and denoised with CaIMan^48^.

#### Intrahippocampal kainic acid mouse model of chronical temporal lobe epilepsy

##### Mice

Male transgenic mice were obtained from The Jackson Laboratory (Nestin-CreER^T2^:016261; ROSA26-CAG-stop^flox^-tdTomato Ai9:007909) to establish a local breeding colony on a C57BL/6J background. Mice were housed in the vivarium on a 12h light/dark cycle, were housed 3-5 mice per cage, and had access food and water ad libitum. Mice were housed individually during video-EEG monitoring following kainic acid injection. Mature male and female mice (>8 weeks of age) were used for all experiments.

##### Experimental timeline

Kainic Acid (KA) was injected into the ventral hippocampus to induce the epilepsy model and rAAV(Syn-GCaMP6f) was injected into the dorsal dentate gyrus ipsilateral to KA injection. Shortly after, a chronic imaging window was implanted over dorsal dentate and an LFP electrode was inserted adjacent to site of KA injection. Following recovery from injection of KA, mice were placed in video-EEG enabled housing where LFP and behavioral activity were continuously recorded to monitor ictogenesis. Three weeks post-KA injection, the video-EEG verified TLE mice were habituated to being head fixed under the two-photon microscope and concurrent Ca2+ imaging and LFP recording was performed. Detailed surgical procedures are reported in^31^.

##### Two-photon imaging

Two-photon imaging of dentate gyrus granule cells was performed using the same set up as^49^, at 4 images/second. Approximately 50-100 mW of laser power under the objective was used for excitation (Ti:Sapphire laser, (Chameleon Ultra II, Coherent) tuned to 920 nm), with adjustments in power levels to accommodate varying window clarity. To optimize light transmission, the angle of the mouse’s head was adjusted using two goniometers (Edmund Optics, +/-10-degree range) such that the imaging window was parallel to the objective. A piezoelectric crystal was coupled to the objective (Nikon 40X NIR water-immersion, 0.8 NA, 3.5mm WD), allowing for rapid displacement of the imaging plane in the z-dimension. OASIS^50^ was used for denoising. While not being imaged, mice were routinely monitored for interictal and seizure events using a custom continuous video-EEG system previously described^6^. Healthy control mouse data was obtained from^49^, included two-photon recording of granule cells from approximately the same location in dentate gyrus as in epileptic mice.

### EFFECTIVE CONNECTIVITY MODELING

#### Modeling neural dynamics in healthy and pathological brains

##### Zebrafish

The start of the preseizure state was defined as one minute after PTZ bath application. Seizure initiation was defined as three standard deviations above population mean calcium signal during baseline (pre-PTZ) and termination was defined as when the population mean signal dropped below this threshold. Correlation matrix analysis^51^ was deployed to quantify network synchronization. Modeling in zebrafish was performed on baseline control dynamics first, and the best fit model was used as the initial parameter matrix for learning preseizure dynamics. Community detection using Leiden algorithm^52^ on learned parameter matrix was used to identify communities with numerous intra-group connections.

##### Mice

For both control and chronically epileptic mice, calcium dynamics were acquired during a 30-minute epoch. Calcium recordings included artifacts from running and grooming that resulted in prolonged hyper-synchronous events. Therefore, a continuous 3-5 minute time window lacking such events was identified for each mouse and used to model calcium dynamics.

#### Chaotic Recurrent Neural Networks

Models of cell-cell effective communication were built using chaotic recurrent neural networks (RNN)^21^. Each neuron experimentally imaged was represented as a node in the network and edges represent the effective (causal) influence between pairs of nodes. The governing dynamics of the chaotic RNN are:

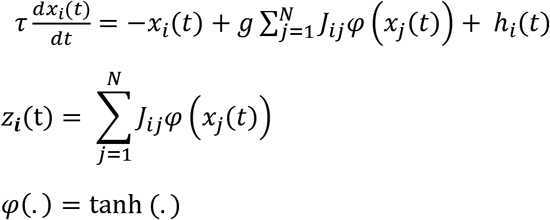

*x*_*i*_(*t*) is the inferred intracellular current of node *i*. *φ*(*x*_*i*_(*t*)) represents firing rate. *z*_*i*_(t) is the estimated calcium signal. *τ* is the time constant of the system (zebrafish: 1.5 s; mouse: 0.625 s). ℎ_*i*_(*t*) is uncorrelated white noise sampled from a normal distribution with mean 0 and standard deviation 0.05 (zebrafish) / 0.005 (mouse). *g* is the gain parameter that determines whether there will be chaos in the system (g<1: stable equilibrium; g>1: chaos). We used g=1.25 for both zebrafish and mouse^25,36^. Lastly, *J* is the effective connectivity matrix that represents the parameters of the model. At initiation of learning baseline control dynamics in zebrafish and learning both control and preseizure dynamics in mice, the values of *J* that were not removed after applying a sparsity mask (zebrafish: 0.10; mouse: 0.40) were sampled from a normal distribution with mean 0 and standard deviation 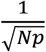. *N* is the number of nodes and *p* is the sparsity. The models contain *p* * *N* * (*N* − 1) parameters. Dynamics were solved using Euler’s method (dt=0.25 s).

#### Reproducing experimental calcium data using FORCE learning

The effective connectivity parameter matrix was optimized to reproduce the experimental calcium data through FORCE learning^22^. This was done through recursive least-squares optimization where at each time point an error signal *e*_*i*_(*t*) = *z*_*i*_(*t*) − *f*_*i*_(*t*) is calculated between node output *z*_*i*_(*t*) and experimental calcium trace *f*_*i*_(*t*). Given this error, the traditional learning rule is as follows:

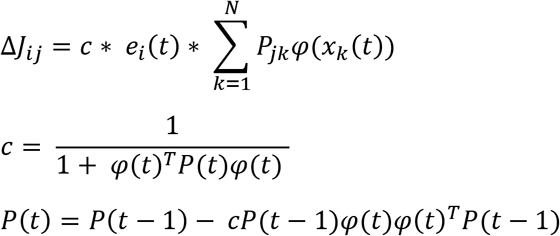

Where *c* is the effective learning rate used for stability and *P(t)* is updated in a recursive fashion at every time point. At initiation, *P(0)* was set to the identify matrix. Learning was performed for 500 epochs for all zebrafish and mouse models, where one epoch is defined as one run of the optimization routine for the duration of the experimental trial. Convergence was assessed by tracking the mean squared reconstruction error between unit activity and experimental data.

#### Biologically constrained FORCE using zebrafish structural connectome

The zebrafish structural connectome^23^ was incorporated into zebrafish FORCE learning as a biological constraint. The structural connectome is a weighted undirected graph between 30 distinct subregions occupying the telencephalon, diencephalon, mesencephalon, and rhombencephalon. The modified weight-update step is:

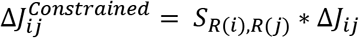

where *i,j* are individual nodes (neurons) and *S*_*R(i),R(j)*_ is the structural connectivity between subregions *R(i)* and *R(j)*.

### DATA ANALYSIS AND QUANTIFICATION

#### Identifying traditional hub neurons

The optimized effective connectivity parameter matrix was binarized by converting the top 10% of edges to a 1 and converting the remaining edges to a 0. The outgoing and incoming degree was calculated for each neuron. Neurons above the 90^th^ percentile outgoing (incoming) degree score were marked as outgoing (incoming) hubs.

#### Force-directed graphs

The effective connectivity matrix was visualized using the Barnes-Hut N-body simulation algorithm^53^. Here, each node can be thought of as a repelling particle and the edges between nodes are modeled as attractive springs. The objective of the algorithm is to find an optimal force-directed spatial configuration such that (i.e. net force = 0). The Barnes-Hut algorithm was applied to the binarized parameter matrix after thresholding for strongest connections (see *Identifying traditional hub neurons*) and used to visualize incoming and outgoing hubs connections in zebrafish.

#### Perturbation simulations and trajectory deviation

To simulate the effect of perturbation on network dynamics, a single hub neuron was ‘current clamped’ to achieve maximum firing rate (500ms step current). The current clamp perturbation was initiated at 0.2 normalized time, consistent for all zebrafish and mouse models. Synthetic calcium traces were then generated for each neuron before and after perturbation.

The Euclidean distance *d* between the mean calcium signal of network with perturbation and with the mean calcium signal of network without perturbation was calculated.

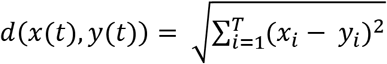

To compare simulations across control and epileptic networks, trajectory deviation *TD* was calculated as the Euclidean distance normalized by remaining time points left after start of perturbation *t*_*o*_.

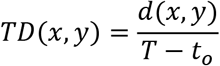

#### Quantifying motif conductance of hubs using local higher-order clustering

Motif-based approximate Personalized PageRank (MAPPR) algorithm^24^ was used to identify an optimal higher-order cluster (i.e. minimum motif conductance) surrounding a hub neuron. Given a motif *M*, MAPPR has three key steps.

##### Constructing motif weighted graph W

The input graph, taken to be the binarized parameter matrix after thresholding for strongest connections (see *Identifying traditional hub neurons*), is transformed into a weighted graph *W* where the weight depends on *M*. Specifically, *W*_*ij*_ is the number of instances of *M* containing nodes *i* and *j*.

##### Compute the approximate personalized PageRank vector

The personalized PageRank (PPR) vector represents the stationary distribution of a modified random walk seeded on a hub node *u.* At each step of the random walk, the random walker is ‘teleported’ back to the specified seed node with probability 1 – α, where α was set to 0.98. The stationary distribution of this process for a seed node *u* (the PPR vector *p*_*u*_), will have larger values for nodes “close” to *u*. The stationary distribution is the solution to the following system of equations:

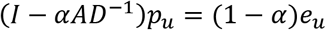

Where *I* is the identify matrix, *A* is the adjacency matrix, *D* is the diagonal degree matrix, and *e*_*u*_ is the vector of all 0’s except for a 1 in position *u*. The PPR vector *p*_*u*_ can be approximated via 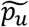 with accuracy *ε* = 0.0001 such that

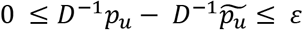

##### Identify higher-order cluster as set with minimal motif conductance

A sweep procedure is used given approximated APPR vector 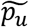. Nodes are sorted by descending value of the vector 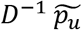 and are incorporated one at a time into a growing set X. After a node added to the set, the motif conductance is calculated.

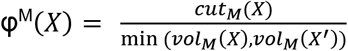

where *vol*_*M*_(*X*) is the number of motif endpoints in X, , *vol*_*M*_(*X*′) is the number of motif endpoints for nodes not in X, and *cut*_*M*_(*X*) is the number of instances of *M* that have at least one end point in X. The set with the smallest motif conductance is deemed optimal and returned. For traditional edge-based clustering, step one is skipped, and the edge conductance score is quantified as above but replacing motif instance *M* with simple edges. Only clusters with at least 5 nodes were considered for analysis.

#### Statistical analysis

Statistics analysis was performed using python *scipy.stats* package. Comparisons involving more than two groups were p-value adjusted (“p-adjusted”) using Bonferroni correction. p-values are reported in all figure subpanels except when p<0.001. Statistical significance was set at 0.05. No data points were left out.

## Acknowledgements

The authors would like to thank Michael Kunst for access to zebrafish structural connectivity data, members of the Soltesz Lab and Maxwell Collard for helpful discussions. D.H. was supported by Stanford Interdisciplinary Graduate Fellowship in coordination with Stanford Wu Tsai Neurosciences Institute (Anonymous Donor). M.L-B. was supported by NIH K99MH112840. Z.L. was supported by 1F31NS120783-01. S.C.B. was supported by NIH R01NS096976 and NIH R01NS103139. I.S. and A.L. were supported by NIH R01NS094668 and NIH U19NS104590. Models were built on the Texas Advanced Computing Center (TACC) Frontera system.

## Author Contributions

D.H., M.L-B., S.C.B., J.L., A.L., K.D., I.S. designed research; M.L-B., F.S., Z.L. collected data; D.H. performed modeling and simulations; D.H, I.R., I.S. analyzed data; D.H., I.S. wrote manuscript.

## Declaration of Interests

The authors declare no competing interests related to this work.

## Code and data availability

All code was written in python3 and is available at https://github.com/dhadjia1/soltesz-lab-epilepsy-modeling. Data are available upon reasonable request.

## Notes

### Competing Interest Statement

The authors have declared no competing interest.

### Summary of Updates

Updated title; focused on single cell control of epileptic circuits

## References

1. Bullmore, E. & Sporns, O. Complex brain networks: graph theoretical analysis of structural and functional systems. 13 (2009).

2. Fornito, A., Zalesky, A. & Breakspear, M. The connectomics of brain disorders. Nature Reviews Neuroscience 16, 159–172 (2015).

3. Kramer, M. A. & Cash, S. S. Epilepsy as a Disorder of Cortical Network Organization. Neuroscientist 18, 360–372 (2012).

4. Jiruska, P. et al. Synchronization and desynchronization in epilepsy: controversies and hypotheses. The Journal of Physiology 591, 787–797 (2013).

5. Löscher, W. & Schmidt, D. Modern antiepileptic drug development has failed to deliver: ways out of the current dilemma. Epilepsia 52, 657–678 (2011).

6. Krook-Magnuson, E., Armstrong, C., Oijala, M. & Soltesz, I. On-demand optogenetic control of spontaneous seizures in temporal lobe epilepsy. Nature Communications 4, 1376 (2013).

7. Bui, A. D. et al. Dentate gyrus mossy cells control spontaneous convulsive seizures and spatial memory. Science 359, 787–790 (2018).

8. Krook-Magnuson, E. & Soltesz, I. Beyond the hammer and the scalpel: selective circuit control for the epilepsies. Nature Neuroscience 18, 331–338 (2015).

9. Krook-Magnuson, E. et al. In vivo evaluation of the dentate gate theory in epilepsy. J Physiol 593, 2379–2388 (2015).

10. Barabási, A.-L. & Albert, R. Emergence of Scaling in Random Networks. Science 286, 509–512 (1999).

11. Morgan, R. J. & Soltesz, I. Nonrandom connectivity of the epileptic dentate gyrus predicts a major role for neuronal hubs in seizures. Proceedings of the National Academy of Sciences 105, 6179–6184 (2008).

12. Bonifazi, P. et al. GABAergic hub neurons orchestrate synchrony in developing hippocampal networks. Science 326, 1419–1424 (2009).

13. Bocchio, M. et al. Hippocampal hub neurons maintain distinct connectivity throughout their lifetime. Nature Communications 11, 4559 (2020).

14. Benson, A. R., Gleich, D. F. & Leskovec, J. Higher-order organization of complex networks. Science 353, 163–166 (2016).

15. Sporns, O. & Kötter, R. Motifs in Brain Networks. PLoS Biol 2, (2004).

16. Lovett-Barron, M. et al. Ancestral Circuits for the Coordinated Modulation of Brain State. Cell 171, 1411–1423.e17 (2017).

17. Liu, J. & Baraban, S. C. Network Properties Revealed during Multi-Scale Calcium Imaging of Seizure Activity in Zebrafish. eNeuro 6, (2019).

18. Burrows, D. R. W. et al. Imaging epilepsy in larval zebrafish. Eur J Paediatr Neurol 24, 70–80 (2020).

19. Baraban, S. C., Dinday, M. T. & Hortopan, G. A. Drug screening in Scn1a zebrafish mutant identifies clemizole as a potential Dravet syndrome treatment. Nat Commun 4, 2410 (2013).

20. Baraban, S. C., Taylor, M. R., Castro, P. A. & Baier, H. Pentylenetetrazole induced changes in zebrafish behavior, neural activity and c-fos expression. Neuroscience 131, 759–768 (2005).

21. Sompolinsky, H., Crisanti, A. & Sommers, H. J. Chaos in Random Neural Networks. Phys. Rev. Lett. 61, 259–262 (1988).

22. Sussillo, D. & Abbott, L. F. Generating Coherent Patterns of Activity from Chaotic Neural Networks. Neuron 63, 544–557 (2009).

23. Kunst, M. et al. A Cellular-Resolution Atlas of the Larval Zebrafish Brain. Neuron 103, 21–38.e5 (2019).

24. Yin, H., Benson, A. R., Leskovec, J. & Gleich, D. F. Local Higher-Order Graph Clustering. in Proceedings of the 23rd ACM SIGKDD International Conference on Knowledge Discovery and Data Mining 555–564 (Association for Computing Machinery, 2017). doi:10.1145/3097983.3098069.

25. Andalman, A. S. et al. Neuronal Dynamics Regulating Brain and Behavioral State Transitions. Cell 177, 970–985.e20 (2019).

26. Song, S., Sjöström, P. J., Reigl, M., Nelson, S. & Chklovskii, D. B. Highly Nonrandom Features of Synaptic Connectivity in Local Cortical Circuits. PLoS Biol 3, (2005).

27. Huang, R. Q. et al. Pentylenetetrazole-induced inhibition of recombinant gamma-aminobutyric acid type A (GABA(A)) receptors: mechanism and site of action. J. Pharmacol. Exp. Ther. 298, 986–995 (2001).

28. Sadeh, S. & Clopath, C. Theory of neuronal perturbome in cortical networks. PNAS 117, 26966–26976 (2020).

29. Albert, R., Jeong, H. & Barabási, A.-L. Error and attack tolerance of complex networks. Nature 406, 378–382 (2000).

30. Muldoon, S. F., Soltesz, I. & Cossart, R. Spatially clustered neuronal assemblies comprise the microstructure of synchrony in chronically epileptic networks. PNAS 110, 3567–3572 (2013).

31. Sparks, F., Liao, Z., Li, W., Soltesz, I. & Losonczy, A. Adult-born granule cells support pathological microcircuits in the chronically epileptic dentate gyrus. http://biorxiv.org/lookup/doi/10.1101/2020.05.01.072173 (2020) doi:110.1101/2020.05.01.072173.

32. Gotman, J. & Pittau, F. Combining EEG and fMRI in the study of epileptic discharges. Epilepsia 52 Suppl 4, 38–42 (2011).

33. Farrell, J. S., Nguyen, Q.-A. & Soltesz, I. Resolving the Micro-Macro Disconnect to Address Core Features of Seizure Networks. Neuron 101, 1016–1028 (2019).

34. Soltesz, I. & Losonczy, A. CA1 pyramidal cell diversity enabling parallel information processing in the hippocampus. Nature Neuroscience 21, 484–493 (2018).

35. Rosch, R. E., Hunter, P. R., Baldeweg, T., Friston, K. J. & Meyer, M. P. Calcium imaging and dynamic causal modelling reveal brain-wide changes in effective connectivity and synaptic dynamics during epileptic seizures. PLOS Computational Biology 14, e1006375 (2018).

36. Rajan, K., Harvey, C. D. & Tank, D. W. Recurrent Network models of sequence generation and memory. Neuron 90, 128–142 (2016).

37. Li, N., Daie, K., Svoboda, K. & Druckmann, S. Robust neuronal dynamics in premotor cortex during motor planning. Nature 532, 459–464 (2016).

38. Ahrens, M. B., Orger, M. B., Robson, D. N., Li, J. M. & Keller, P. J. Whole-brain functional imaging at cellular resolution using light-sheet microscopy. Nature Methods 10, 413–420 (2013).

39. Betzel, R. F. Organizing principles of whole-brain functional connectivity in zebrafish larvae. Network Neuroscience 4, 234–256 (2019).

40. Kobayashi, M. & Buckmaster, P. S. Reduced Inhibition of Dentate Granule Cells in a Model of Temporal Lobe Epilepsy. J. Neurosci. 23, 2440–2452 (2003).

41. Korn, S. J., Giacchino, J. L., Chamberlin, N. L. & Dingledine, R. Epileptiform burst activity induced by potassium in the hippocampus and its regulation by GABA-mediated inhibition. J. Neurophysiol. 57, 325–340 (1987).

42. Isaacson, J. S. & Scanziani, M. How inhibition shapes cortical activity. Neuron 72, 231–243 (2011).

43. Murphy, B. K. & Miller, K. D. Balanced Amplification: A New Mechanism of Selective Amplification of Neural Activity Patterns. Neuron 61, 635–648 (2009).

44. Chang, W.-C. et al. Loss of neuronal network resilience precedes seizures and determines the ictogenic nature of interictal synaptic perturbations. Nature Neuroscience 21, 1742–1752 (2018).

45. Ponce-Alvarez, A., Jouary, A., Privat, M., Deco, G. & Sumbre, G. Whole-Brain Neuronal Activity Displays Crackling Noise Dynamics. Neuron 100, 1446–1459.e6 (2018).

46. Shew, W. L., Yang, H., Petermann, T., Roy, R. & Plenz, D. Neuronal Avalanches Imply Maximum Dynamic Range in Cortical Networks at Criticality. J. Neurosci. 29, 15595–15600 (2009).

47. Freeman, J. et al. Mapping brain activity at scale with cluster computing. Nature Methods 11, 941–950 (2014).

48. Giovannucci, A. et al. CaImAn an open source tool for scalable calcium imaging data analysis. eLife 8, e38173 (2019).

49. Danielson, N. B. et al. Distinct Contribution of Adult-Born Hippocampal Granule Cells to Context Encoding. Neuron 90, 101–112 (2016).

50. Friedrich, J., Zhou, P. & Paninski, L. Fast online deconvolution of calcium imaging data. PLOS Computational Biology 13, e1005423 (2017).

51. Li, X. et al. Synchronization measurement of multiple neuronal populations. Journal of Neurophysiology 98, 3341–3348 (2007).

52. Traag, V. A., Waltman, L. & van Eck, N. J. From Louvain to Leiden: guaranteeing well-connected communities. Scientific Reports 9, 5233 (2019).

53. Barnes, J. & Hut, P. A hierarchical O(N log N) force-calculation algorithm. Nature 324, 446–449 (1986).

