## Supplementary figure legends and supplementary figures for "Maximally selective single cell target for circuit control in epilepsy"

**Supplementary Figure 1: Macroscale anatomical subnetworks emerge from models constrained by zebrafish structural connectome**

Community detection using the Leiden algorithm partitioned the graph into multiple communities. The algorithm optimizes the partition such that each community has dense connections between its constituents while there are sparse connections between nodes in different communities. Community detection was performed on FORCE optimized networks with **(A)** and without **(B)** the structural constraint, showing that structurally constrained optimization partitioned the network into relevant macroscale subnetworks.

**Supplementary Figure 2: Additional validation of FORCE optimized chaotic recurrent neural networks**

**(A,B):** Models were trained in the absence of controlled external manipulation Despite this, optimized models are primarily driven by synaptic inputs and not noise for both baseline (A) and preseizure (B) networks.

**C:** Scatter plot of total inhibitory output from neurons in baseline (x-axis) and preseizure (y-axis) networks. Total inhibitory output was taken to be the sum of outgoing negative weights. Linear regression analysis validates that inhibition decreases during bath wash-in of PTZ for all modeled fish, as the slope parameter *a* of the dashed lines ($Inh^{preseizure} = a*Inh^{baseline} + b$) is less than 1.

**Supplementary Figure 3: Additional characterization of network response to single-cell perturbation in zebrafish cohort**

**(A,B):** Average calcium signal of nodes receiving strong excitatory inputs from a single outgoing that underwent (t=0) simulated depolarizing current injection in the baseline (A) and preseizure (B) network.

**C:** Trajectory deviation of outgoing, incoming, and non-hub neurons in baseline network. Perturbation of a single outgoing hub had significantly higher effect on global network dynamics that incoming hubs and non-hubs (one-sided Mann-Whitney U-test. p<0.001 after Bonferroni correction). Perturbation of non-hubs had significantly higher effect on global dynamics than incoming hubs (one-sided Mann-Whitney U-test. p<0.001 after Bonferroni correction).

**D:** Trajectory deviation of preseizure network showing similar results to A (one-sided Mann-Whitney U-test. p <0.001 after Bonferroni correction).

**Supplementary Figure 4: Disconnecting superhubs more effectively stabilized network response to perturbation compared to disconnecting random hubs or low-conductance hubs**

**A:** Comparison of change in signal variance before and after perturbation after disconnecting superhubs and equal number of hubs randomly. Disconnecting superhubs was more effective in stabilizing networks to perturbation than disconnecting hubs randomly (two-sided Wilcoxon signed-rank test, p<0.001).

**B:** Similar to *A*, disconnecting superhubs hubs was more effective in stabilizing networks to perturbation than disconnecting equal number of low-conductance hubs (two-sided Wilcoxon signed-rank test, p<0.001).

**Supplementary Figure 5: Additional characterization of network response to single hub perturbation in mouse cohort**

**A:** Violin plots of trajectory deviation in response to single outgoing an incoming hub perturbation. Perturbation of outgoing hubs in control dentate network has more significant influence on global network dynamics than incoming hubs (one-sided Mann-Whitney U-test, p<0.001).

**B:** Perturbation of outgoing hubs in chronically epileptic dentate network has more significant influence on global network dynamics than incoming hubs (one-sided Mann-Whitney U-test, p<0.001).

**C:** Trajectory deviation of fully connected and disconnected chronically epileptic dentate network quantified from response to perturbation of individual outgoing hubs. There is less change to global network dynamics when perturbing outgoing hubs in the disconnected network compared to the fully connected chronically epileptic dentate network (one-sided Wilcoxon signed-rank test, $p<0.001$).

**D**: Comparison of change in signal variance before and after perturbation after disconnecting superhubs and equal number of hubs randomly. Disconnecting superhubs was more effective in stabilizing chronically epileptic dentate networks to perturbation than disconnecting hubs randomly (two-sided Wilcoxon signed-rank test, $p=0.0077$).

**E:** Similar to *D*, disconnecting superhubs was more effective in stabilizing chronically epileptic dentate networks to perturbation than disconnecting equal number of low-conductance hubs (two-sided Wilcoxon signed-rank test, $p=0.0024$).

### Supplementary Figure 1

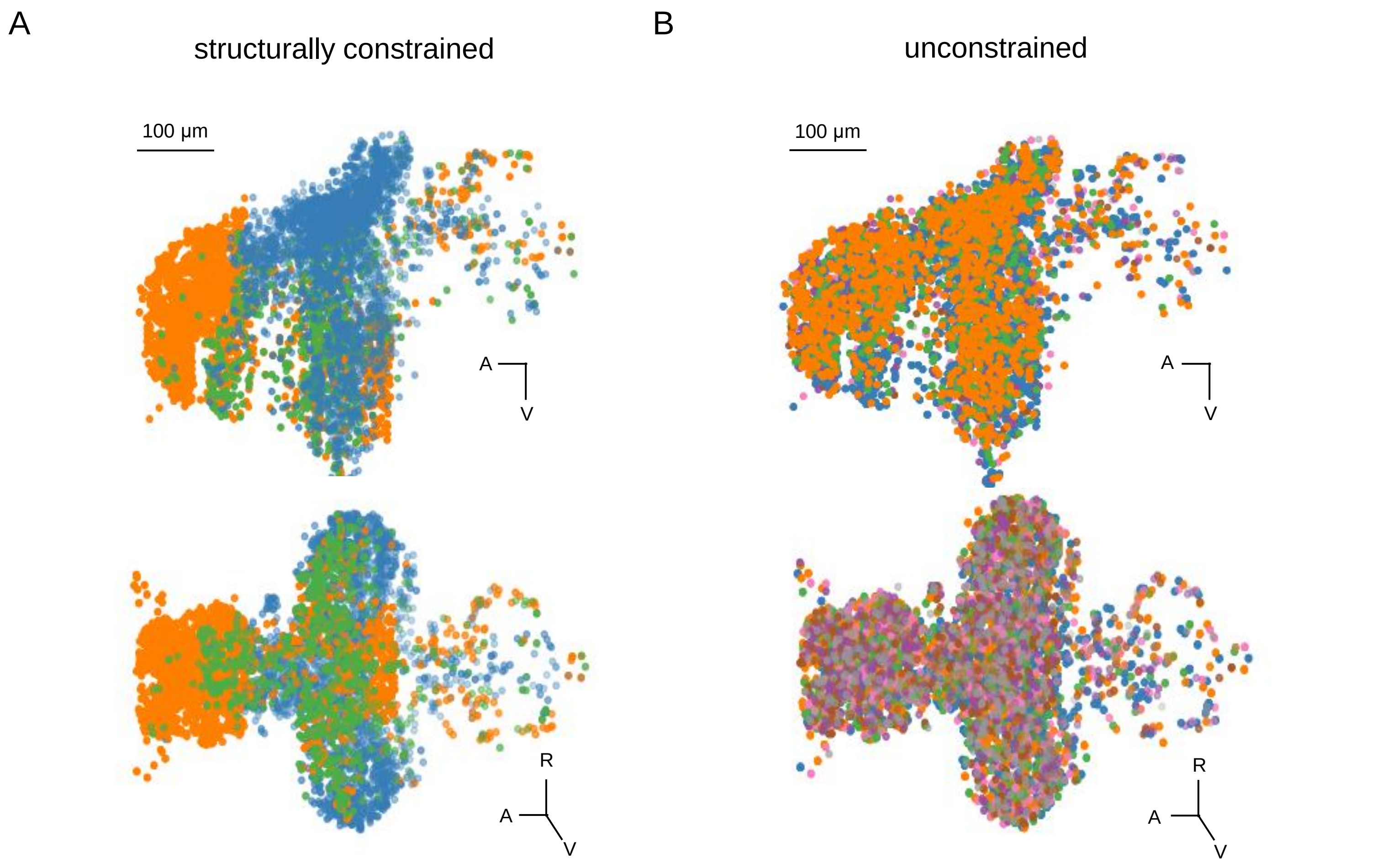

### Supplementary Figure 2

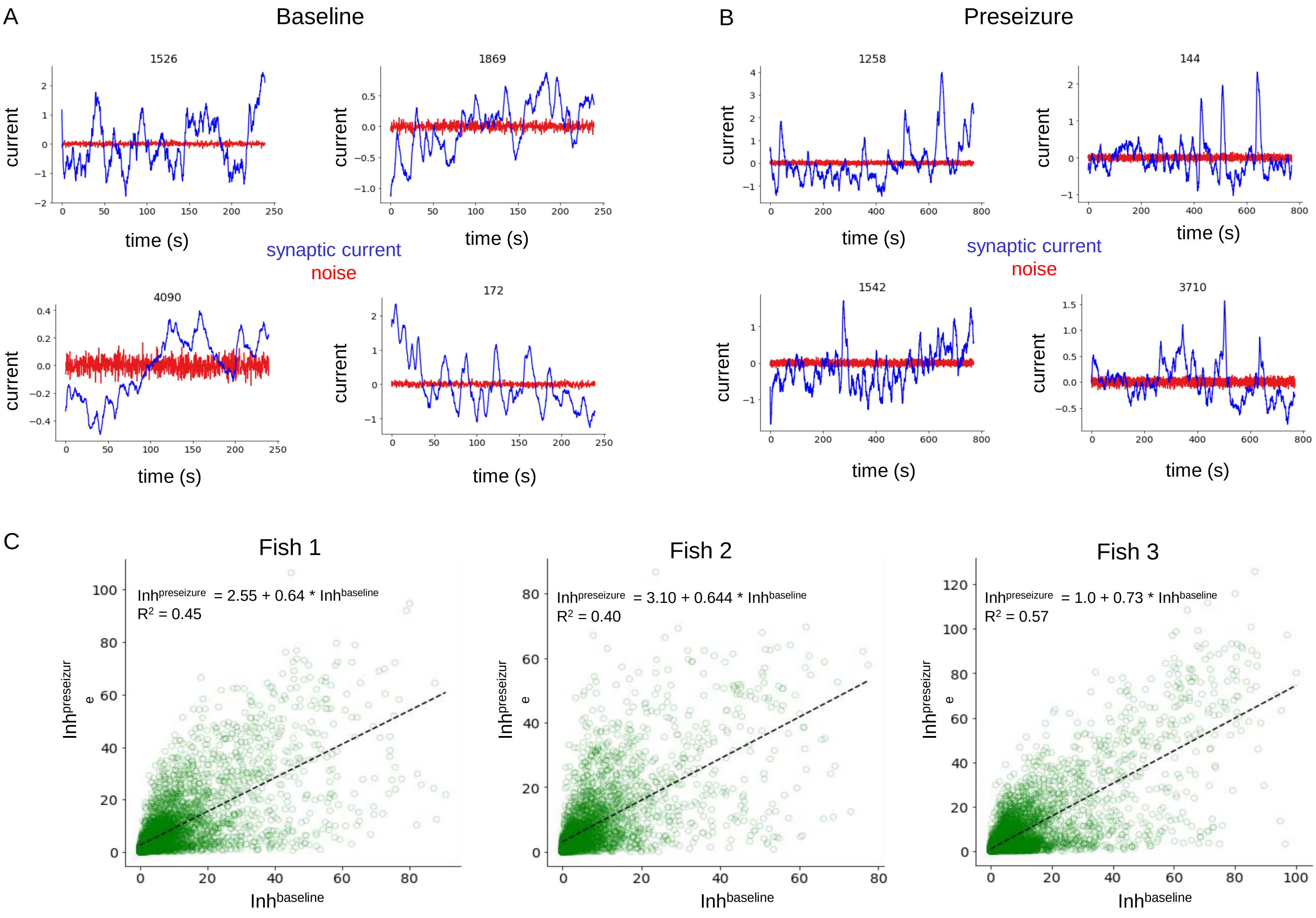

### Supplementary Figure 3

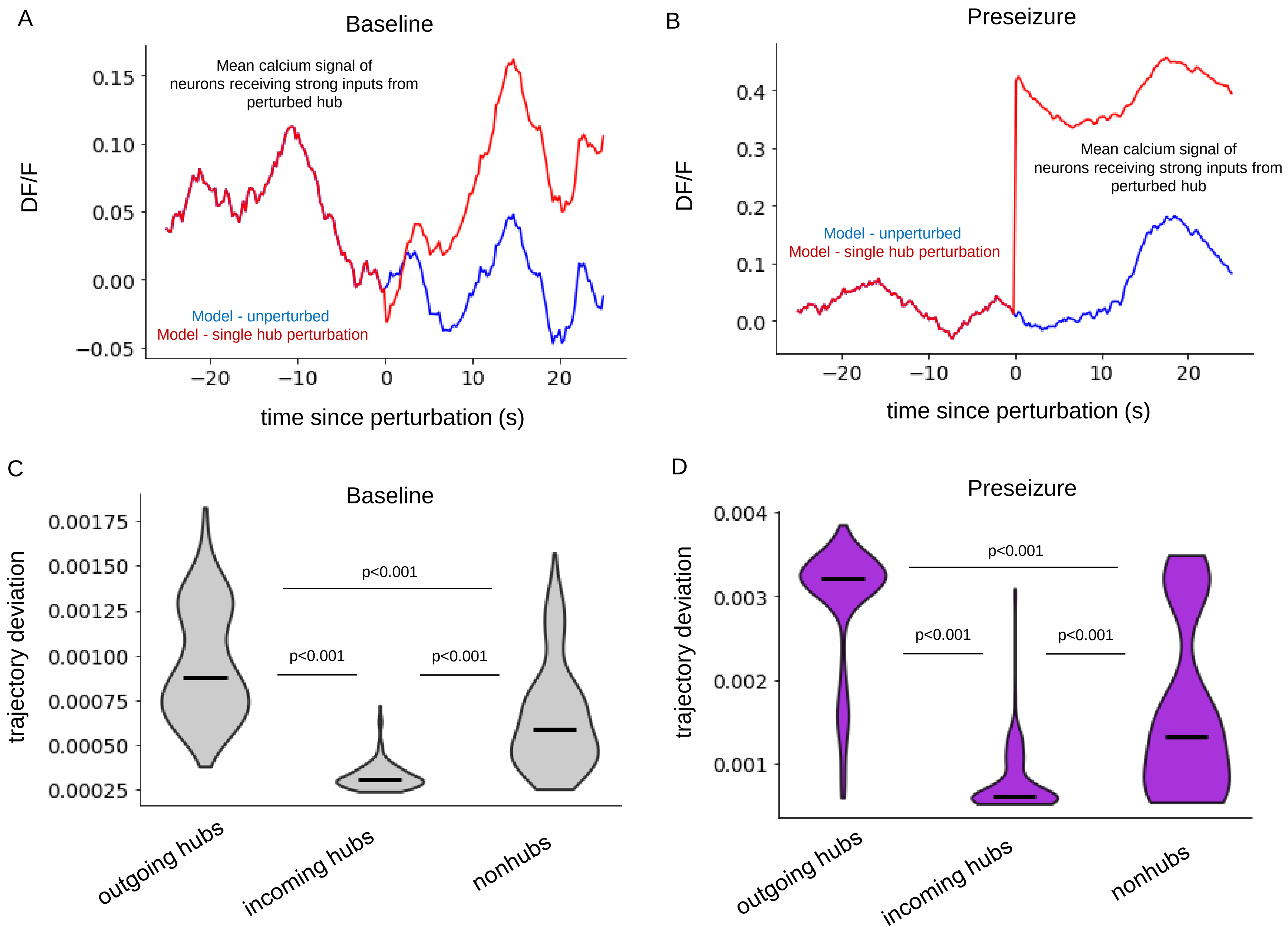

#### Supplementary Figure 4

A

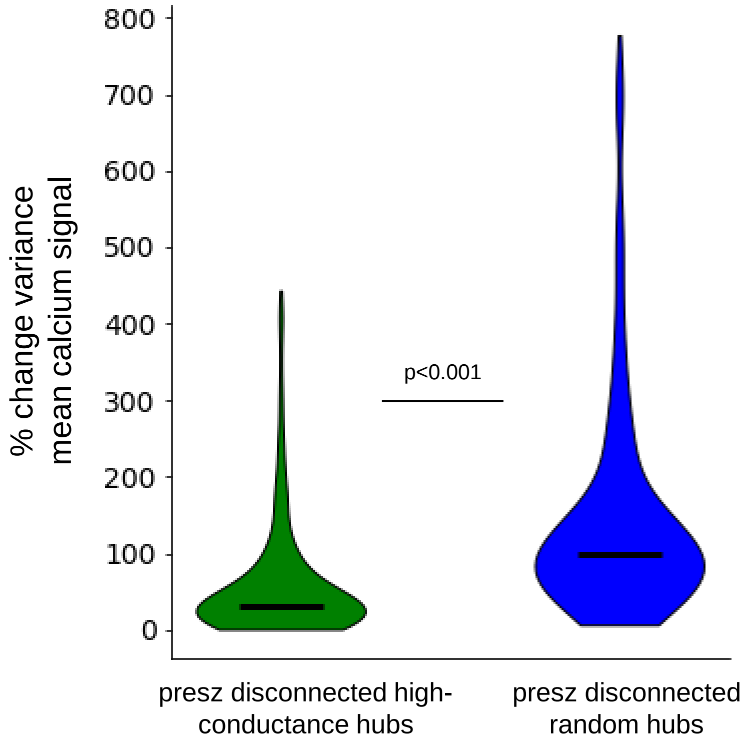


B

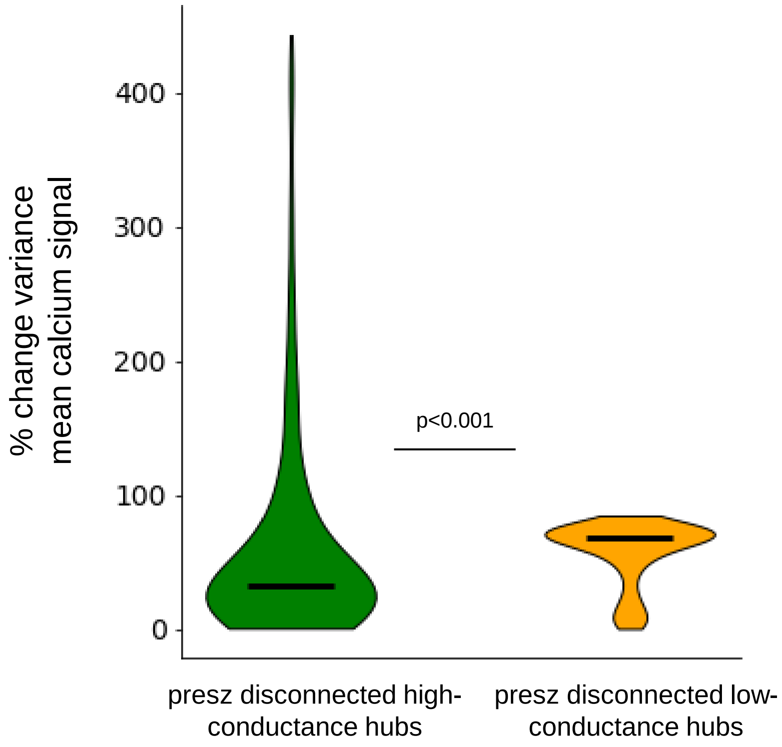

### Supplementary Figure 5

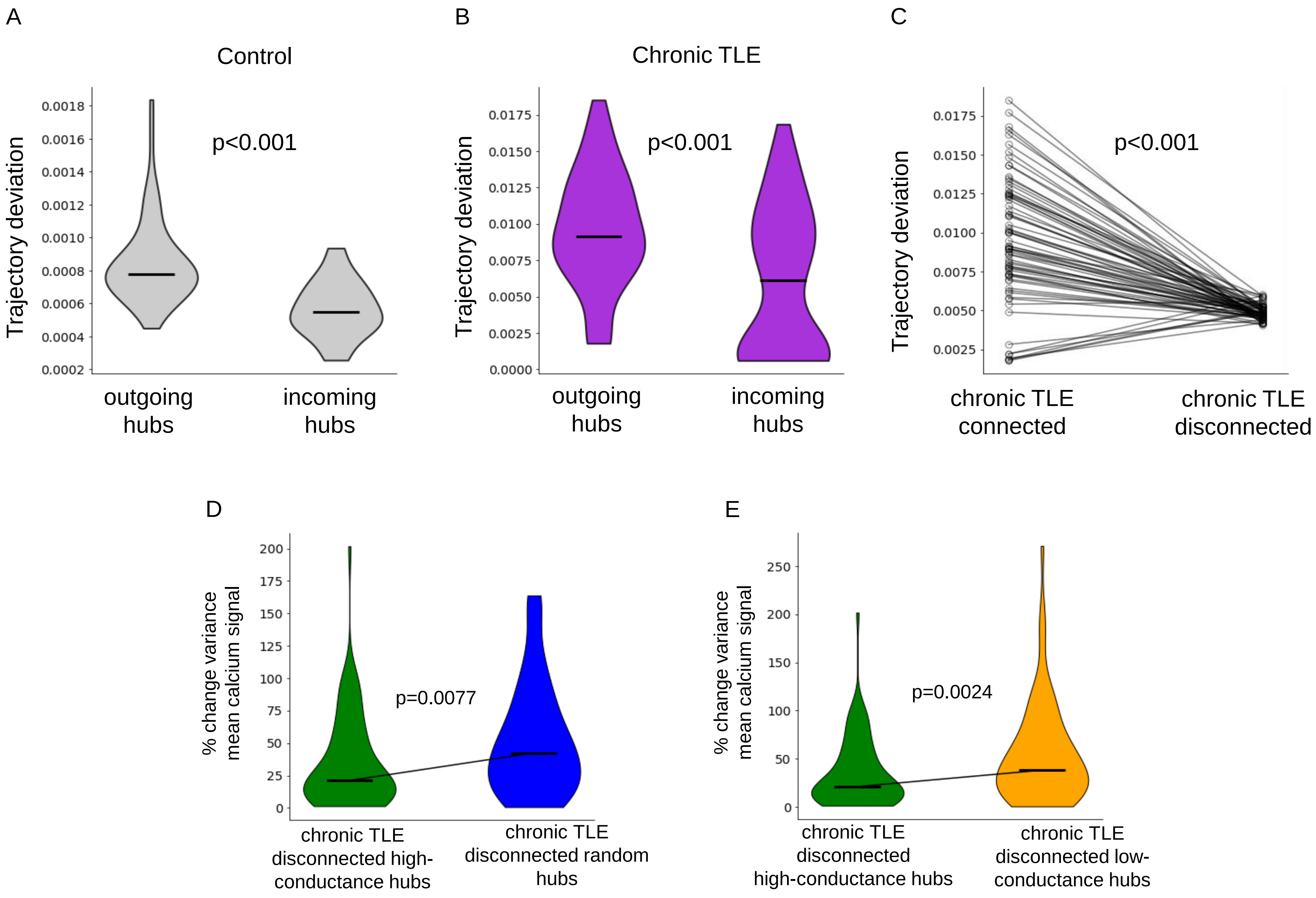
